# Genomic evidence reveals SPA-regulated developmental and metabolic pathways in dark-grown *Arabidopsis* seedlings

**DOI:** 10.1101/2020.02.10.941708

**Authors:** Vinh Ngoc Pham, Inyup Paik, Ute Hoecker, Enamul Huq

**Author notes:** Corresponding author: Enamul Huq, University of Texas at Austin, NHB 2.616, Stop A5000, 100 East 24th St. Austin, TX 78712-1095. **Abbreviations:** COP1: CONSTITUTIVE PHOTOMORPHOGENIC 1; CRL: CULLIN RING E3 LIGASE; DET1: DE-ETIOLATED1; HY5: ELONGATED HYPOCOTYL 5; LEA: LATE EMBRYOGENESIS ABUNDANT PROTEIN GENES; PIF: PHYTOCHROME INTERACTING FACTOR; FPKM: FRAGMENTS PER KILO BASES PER MILLION READS; RNA-Seq: RNA sequencing; SPA: SUPPRESSOR OF PHYA-105.

## Abstract

Photomorphogenesis is repressed in the dark mainly by an E3 ubiquitin ligase complex comprising CONSTITUTIVE PHOTOMORPHOGENIC 1 (COP1) and four homologous proteins called SUPPRESSOR OF PHYA-105 (SPA1-SPA4) in Arabidopsis. This complex induces the ubiquitination and subsequent degradation of positively acting transcription factors (e.g., HY5, HFR1, PAP1 and others) in the dark to repress photomorphogenesis. Genomic evidence showed a large number of genes regulated by COP1 in the dark, of which many are direct targets of HY5. However, the genomic basis for the constitute photomorphogenic phenotype of *spaQ* remains unknown. Here, we show that >7200 genes are differentially expressed in the *spaQ* background compared to wild-type in the dark. Comparison of the RNA Sequencing (RNA-Seq) data between *cop1* and *spaQ* revealed a large overlapping set of genes regulated by the COP1-SPA complex. In addition, many of the genes coordinately regulated by the COP1-SPA complex are also regulated by HY5 directly and indirectly. Taken together, our data reveal that SPA proteins repress photomorphogenesis by controlling gene expression in concert with COP1, likely through regulating the abundance of downstream transcription factors in light signaling pathways. Moreover, SPA proteins may function both in a COP1-dependent and –independent manner in regulating many biological processes and developmental pathways in Arabidopsis.

**Summary statement:** Comparison of transcriptome analyses between *cop1* and *spaQ* mutants reveal overlapping pathways regulated by COP1 and SPAs.

## Introduction

Phenotypic plasticity provides plants with successful strategies to adapt to the prevailing environmental conditions. Upon germination, seedlings undergo two developmental fates depending on the absence or presence of light. In the absence of light, seedlings undergo skotomorphogenesis; the hypocotyls elongate and closed cotyledons and an apical hook protect the shoot apical meristem. By contrast, seedlings display a short hypocotyl, no apical hook, open, expanded and green cotyledons in the presence of light in a process called photomorphogenesis (Bae and Choi 2008, Pham et al. 2018a). It has been proposed that photomorphogenesis is the default pathway for plant development as a series of repressor proteins, CONSTITUTIVE PHOTOMORPHOGENIC/DE-ETIOLATED1/FUSCA (COP/DET/FUS), have been identified that inhibit photomorphogenesis in the dark. Mutants with defects in any of these inhibitors display constitutive photomorphogenic (cop) phenotypes in the dark (Lau and Deng 2012).

The repressor proteins are classified into four groups with overlapping functions. The first one is called CONSTITUTIVE PHOTOMORPHOGENIC1 (*COP1*), which encodes a RING finger protein containing WD40 repeats and a coiled-coil domain (Deng et al. 1992). COP1 functions as a central repressor and an E3 ubiquitin ligase to promote the ubiquitination and degradation of transcription factors that act positively in photomorphogenesis (e.g., HY5/LAF1/HFR1 and others) (Hardtke et al. 2000, Osterlund et al. 2000, Seo et al. 2003, Jang et al. 2005, Yang et al. 2005). The degradation of these positive factors results in repression of photomorphogenesis in the dark. Extensive research also showed that COP1 not only functions as a central regulator in light signaling but also in regulation of flowering time, circadian rhythm, UV-B signaling, shade avoidance response, cold acclimation response and many other pathways (Hoecker 2017).

COP1 also interacts with the four members of the SUPPRESSOR OF PHYA-105 family (SPA1-SPA4) to form tetrameric complexes consisting of two COP1 and two SPA proteins (COP1/SPA complex) (Zhu et al. 2008). Genetic analysis indicates that SPA proteins strongly enhance COP1 function (Saijo et al. 2003, Seo et al. 2003, Ordoñez-Herrera et al. 2015). Finally, the COP1-SPA complex also associates with CULLIN4 to form a CUL4-COP1-SPA complex that functions as a CULLIN RING E3 LIGASE (CRL) to promote degradation of the positively acting transcription factors in the dark to repress photomorphogenesis (Chen et al. 2010). On the other hand, CUL4^COP1-SPA^ complex can rapidly trigger early ubiquitin-mediated degradation of PHYTOCHROME INTERACTING FACTOR 1(PIF1) to initiate light-induced seed germination (Zhu et al. 2015, Paik et al. 2019). This suggests a unique dual role of CUL4^COP1-SPA^ complex in dark and in early light induced photomorphogenesis.

The COP1/SPA complex is inhibited by light, thus allowing the substrate transcription factors such as HY5 to accumulate in light and induce photomorphogenesis. Light-activated photoreceptors, cryptochromes and phytochromes, directly interact with the COP1/SPA complex and inactivate the complex through multiple mechanisms. COP1/SPA is thereby inactivated through multiple mechanisms, including the nuclear exclusion of COP1, the dissociation of the COP1/SPA complex and the degradation of SPA proteins (Hoecker 2017, Podolec and Ulm 2018). A recent study also showed a non-canonical function of COP1-SPA by inhibiting BIN2 kinase to stabilize phytochrome-interacting factor 3 (PIF) in the dark, which illustrates the dynamic regulatory function of the COP1-SPA complex (Ling et al. 2017, Pham et al. 2018a).

*SPA1* was first identified from a genetic screening as a suppressor of a weak *phyA-105* mutant (Hoecker et al. 1998, Hoecker et al. 1999). Since then, SPA proteins have been shown to play an important role as the core member of the COP1-SPA E3 ligase complex, in regulating photomorphogenesis, together with phytochromes and cryptochromes. Unlike *cop1* null mutants, quadruple *spa* mutants (*spaQ*) are viable beyond seedling stage but exhibit strikingly dwarfed growth (Laubinger et al. 2004, Ordoñez-Herrera et al. 2015). Like hypomorphic *cop1* mutants, *spaQ* mutant undergoes constitutive photomorphogenesis, including short hypocotyl and open cotyledons in the dark, similar to light-grown seedlings, and moreover, fail to show photoperiodic regulation of flowering time (McNellis et al. 1994, Laubinger et al. 2004, Laubinger et al. 2006). Different *SPAs* exhibit redundant functions but also have partially distinct roles in regulating plant development. *SPA1* and *SPA2* primarily regulate seedling growth and development, while *SPA3* and *SPA4* play major roles in leaf expansion in adult plants (Laubinger and Hoecker 2003, Laubinger et al. 2004, Fittinghoff et al. 2006, Hoecker 2017). SPA proteins contain a ser/thr kinase domain at the N-terminus, a coiled-coil domain in the middle, and four WD-40 repeats at the C-terminus (Hoecker et al. 1999). Although, SPA proteins are known for their essential role in enhancing COP1 E3 ligase function, a COP1-independent role for SPAs has not been shown yet, despite the presence of a kinase-like domain at the N-terminus. We have recently shown that the N-terminal domain of SPA1 possesses a ser/thr kinase activity which directly phosphorylates PIF1 to induce rapid light-induced degradation (Paik et al. 2019). Moreover, the kinase domain is also necessary for seedling etiolation in darkness (Holtkotte et al. 2016, Paik et al. 2019).

The second repressor complex is called the COP9 signalosome (CSN) which consists of eight subunits (CSN1-CSN8), that are highly conserved from plants to humans (Serino and Deng 2003). The CSN is involved in deneddylation/derubylation of the CULLIN Ring family of E3 Ubiquitin ligases (CRLs) (Schwechheimer et al. 2001). Many of the CSN subunit mutants were isolated through their cop phenotype in the dark (Serino and Deng 2003).

The third repressor complex consists of DEETIOLATED1 (DET1), COP10, DNA damage-binding protein 1 (DDB1) and CUL4. Among the COP/DET/FUS repressors, DET1 is the first described nuclear localized protein that has been shown to bind to histone H2B (Pepper et al. 1994, Benvenuto et al. 2002). DET1 inhibits seed germination and photomorphogenesis in the dark by oppositely regulating the abundance of PIFs and HFR1 post-translationally (Dong et al. 2014, Shi et al. 2015). DET1 also forms a complex with COP10, DDB1 and CUL4 to form a CUL4^CDD^ complex that degrades positively acting transcription factors to repress photomorphogenesis in the dark (Schroeder et al. 2002, Chen et al. 2006).

The fourth group of repressors has been identified as PHYTOCHROME INTERACTING FACTORs (PIFs). PIFs (PIF1-PIF8) belong to the basic helix-loop-helix (bHLH) classes of transcription factors that interact directly with the active Pfr form of phytochromes (mainly phyB, but PIF1 and PIF3 interact with both phyA and phyB) (Leivar and Quail 2011, Pham et al. 2018a) and mainly repress photomorphogenesis in the dark (Leivar et al. 2008, Shin et al. 2009). PIFs also interact with COP1 and SPA1, and enhance the substrate recruitment, auto- and trans-ubiquitination activities of COP1 (Xu et al. 2014, Xu et al. 2015, Xu et al. 2017). In contrast, COP1 and SPA1 interact with the phosphorylated forms of PIF1 and induce its degradation in a light-dependent manner to promote seed germination (Zhu et al. 2015, Paik et al. 2019). In addition, we have proposed a modified model for the cop phenotypes, which is not only due to an increased abundance of positively acting transcription factors (e.g., HY5, HFR1 and others), but also due to COP1- and SPA-mediated stabilization of PIFs, and HFR1-mediated inhibition of PIF function in darkness (Pham et al. 2018b).

COP1 functions have been extensively investigated by genomic methods including microarray analyses of various *cop1* mutants (Ma et al. 2002). These analyses showed similar expression profile between dark-grown *cop1* mutants and light-grown wild-type seedlings (Ma et al. 2001, Ma et al. 2002). In these studies, COP1 function has been suggested as the central hub for integration of light, hormone and temperature signaling pathways (Ma et al. 2002, Catalá et al. 2011, Menon et al. 2016, Wang et al. 2016). However, it is still unknown if SPA proteins also function together with COP1 as the integrator among light, hormone, temperature and other biological processes. We have recently reported that a large number of genes are differentially expressed in *spaQ* background in the dark and the kinase activity of SPA1 is necessary for the seedling etiolation phenotype in darkness (Paik et al. 2019). However, this study provides comprehensive genomic analyses on how SPA proteins repress photomorphogenesis in the dark. Gene expression profile comparison between *cop1* and *spaQ* mutants and enriched Gene Ontology analysis suggest that COP1 and SPA proteins share a large overlapping set of gene expression and modulate the actions of key transcriptional regulators in light signaling. This study also provides genomic evidence of how SPA proteins might function in a COP1-dependent and -independent manner to regulate gene expression that drives plant development.

## Materials and methods

### Plant materials and RNA-Sequencing (RNA-Seq) experiment

Total RNAs were extracted from 3-day-old dark-grown Col-0, *cop1-4* and *spaQ* seedlings. Plates were kept in the dark for 3 days at 4°C for stratification. Plates were then exposed to white light for 3hrs at room temperature to promote the germination before being placed in the dark for 21 hrs. Seeds were then treated with far-red light (2000 μmolm^-2^) to inactivate phytochrome activity before putting back into the dark for 2 days. RNAs were extracted in Day 3. Samples from three independent replicates for each genotype were performed.

Total RNAs were extracted using the Sigma spectrum total RNA kit (Sigma Aldrich, St. Louis, MO) according to the manufacturer’s manual. Total RNAs were treated with on-column DNase I (Sigma Aldrich, St. Louis, MO). Quality and integrity of RNA were examined using the Agilent RNA 6000 Pico and Nano Kit (Agilent) with Agilent 2100 Bioanalyzer. Library preparation was performed using NEB Next Ultra™ II RNA Library Prep Kit for Illumina. Libraries from three replicates were sequenced (single read 50) on the HiSeq2500 platform.

### Transcriptomic analyses

All data preprocesses including the quality control, alignment, counting reads were performed using Texas Advanced Computing Center high-performance computing. All plots were generated using the DeSeq2 (Love et al. 2014), ggplot2 (Wickham 2016) and ComplexHeatmap packages in R statistical computing environment (Gu et al. 2016). FastQC (http://www.bioinformatics.babraham.ac.uk/projects/fastqc/) was used to check the quality of the RNA-Seq reads. Alignment to the Arabidopsis genome was performed using Bowtie2 with default parameters (mismatch = 0) (Langmead and Salzberg 2012) and analyzes the mapping results to identify splice junctions between exons using TopHat (https://ccb.jhu.edu/software/tophat/index.shtml) (Trapnell et al. 2012, Kim et al. 2013). The annotation of the *A. thaliana* genome (TAIR10 https://www.arabidopsis.org/) in a GTF file were used for mapping of spliced reads. TopHat scripts used fr-unstranded and –G parameter using Arabidopsis reference GTF file.

Aligned BAM files were created and sorted, indexed and viewed in a genome browser IGV. Total read counts for each sample were performed using HTSeq (Anders et al. 2015) to count the aligned reads to features (http://htseq.readthedocs.io/en/master/index.html). HTSeq returns the counts of each gene for every sample. For these analyses, three biological replicates were analyzed separately. HTSeq output data contain read counts for each biological replicate.

For differential gene expression analysis, we applied both DESeq2 (<u>10.18129/B9.bioc.DESeq2</u>) and Cuffdiff (from Cufflinks) http://cufflinks.cbcb.umd.edu/) (Trapnell et al. 2012). DESeq2 package uses negative binomial generalized linear models to identify the log2 fold change values for different pairwise comparison (e.g., *cop1-4*/WT and *spaQ*/WT). In DESeq2 analysis, DESeqDataSet is provided with an experimental design with all three replicates for Col-0, *cop1-4* and *spaQ* under different light treatments. DESeq2 estimation of the dispersion of counts were done using three biological replicates. On the other hand, the aligned reads from TopHat also were further assembled into transcripts and quantified the expression level in each sample using Cufflinks using a rigorous statistical analysis. Comparison of differentially expressed genes in two conditions were performed by Cuffdiff. The normalized expression level (FPKM; fragments per kilobase per million reads) from RNA-Seq data were performed using Cuffdiff. The differential expressed genes were selected for all the genes with |log2 fold-change| > 1 with adjusted P-value < 0.05. Raw data and processed data for the total read counts of sequencing reads and list of differentially gene expressed in *cop1-4* and *spaQ* can be accessed by Gene Expression Omnibus database (GSE112662).

The matrix of scatter plots and volcano plots for visualization of gene expression in wild-type, *cop1-4* and *spaQ* mutants were quantified by normalized number of fragments per kilo bases per million reads (FPKM) using Cufflinks (Trapnell et al. 2012) and Cummerbund package to transform Cuffdiff data into the R statistical computing environment.

Venny 2.1.0 (http://bioinfogp.cnb.csic.es/tools/venny/) was used to generate all the venn diagram in the figures. Hierarchical clustering heatmap was generated using DESeq2 and R. **D**atabase for **A**nnotation, **V**isualization and **I**ntegrated **D**iscovery (**DAVID**) v6.8 https://david.ncifcrf.gov/ was applied for Gene Ontology (GO) enrichment analysis. Significant enrichment terms were defined and generated bar graphs with lowest p-value and FDR < 0.05.

GO Analysis Toolkit and Database for Agricultural Community (AgriGo http://bioinfo.cau.edu.cn/agriGO/index.php) was applied to generate hieratical graph for *spaQ* regulated genes. For comparison between differential genes expressed in *cop1-4*, *spaQ* and differential genes expressed in *hy5* mutants and compare overlapped genes with HY5 direct target genes, the RNA-Seq raw data from *hy5* mutant and HY5-bound gene list were obtained from a previously study (GSE24974, GSM613465 for mRNASeq_WT and GSM613466 for mRNASeq_*hy5-215*) (Zhang et al. 2011). Original expression profiles of *det1-1* dark-grown seedlings and light-grown wild-type seedlings were obtained from (GSE60835) (Dong et al. 2014).

The list of Arabidopsis transcription factors was obtained from Plant Transcription Factor Database (http://planttfdb.cbi.pku.edu.cn/), for identifying the transcription factors which are differentially expressed in *spaQ* mutant.

### Quantitative RT-PCR assay

For the transcript level of HY5 direct target genes and also genes independently expressed in the *cop1* and *spaQ* mutants, total RNAs were extracted under identical conditions as RNA-Seq. Total RNA treated with DNase I (Sigma Aldrich, St. Louis, MO) was reversed transcribed using M-MLV reverse transcriptase (Thermo Fisher Scientific, Bartlesville, OK). Primers using for qRT-PCR were listed in Table S2. PP2A was used as the internal control. The relative gene expression was calculated using the 2^ΔΔ^CT method.

## Results

### Analysis of differentially expressed genes in *spaQ* compared to wild-type

To understand the genomic basis for the constitutive photomorphogenic phenotypes of *spaQ*, RNA isolated from three independent replicates of wild-type, *cop1-4* and *spaQ* mutant seedling grown in darkness was sequenced as described recently (Paik et al. 2019). An average of 34 million raw reads per sample was obtained from sequencing (Table S1). Among the raw reads, 98% were mapped to the genome, and an average of 85% was uniquely mapped. The quality of reads was then assessed by FastQC. To obtain sample to sample distance, we generated principle component analysis (PCA) and also the heat map of the distance matrix using variance stabilizing transformed count matrix and the regularized logarithm transformed data, respectively (Figs. S1-S2). The raw counts for each sample obtained from HTSeq were then analyzed using DESeq2 package for differential gene expression. Furthermore, we also applied Cuffdiff analysis with the cut-off value for adjusted p-value of 0.05 to identify significantly differentially expressed genes. Global comparison of expression values for each gene in every pairwise comparison have been shown in scatterplot matrix (Fig. S3) and volcano matrix to inspect differentially expressed genes between two samples (Fig. S4). These analyses show that the gene expression profile of *spaQ* is strongly different from wild-type samples.

The significantly differentially expressed genes between wild-type (WT) and *spaQ* seedlings are presented in the MA plot with log2 fold-change of each gene over the mean of normalized counts (Fig. 1A). MA plot derived from DESeq2 shows that most of the genes with higher average normalized counts produce a significant expression call, represented by red dots including both up-regulated genes (positive log2 fold-change values) and down-regulated genes (negative log2 fold-change values).

**Figure 1.**
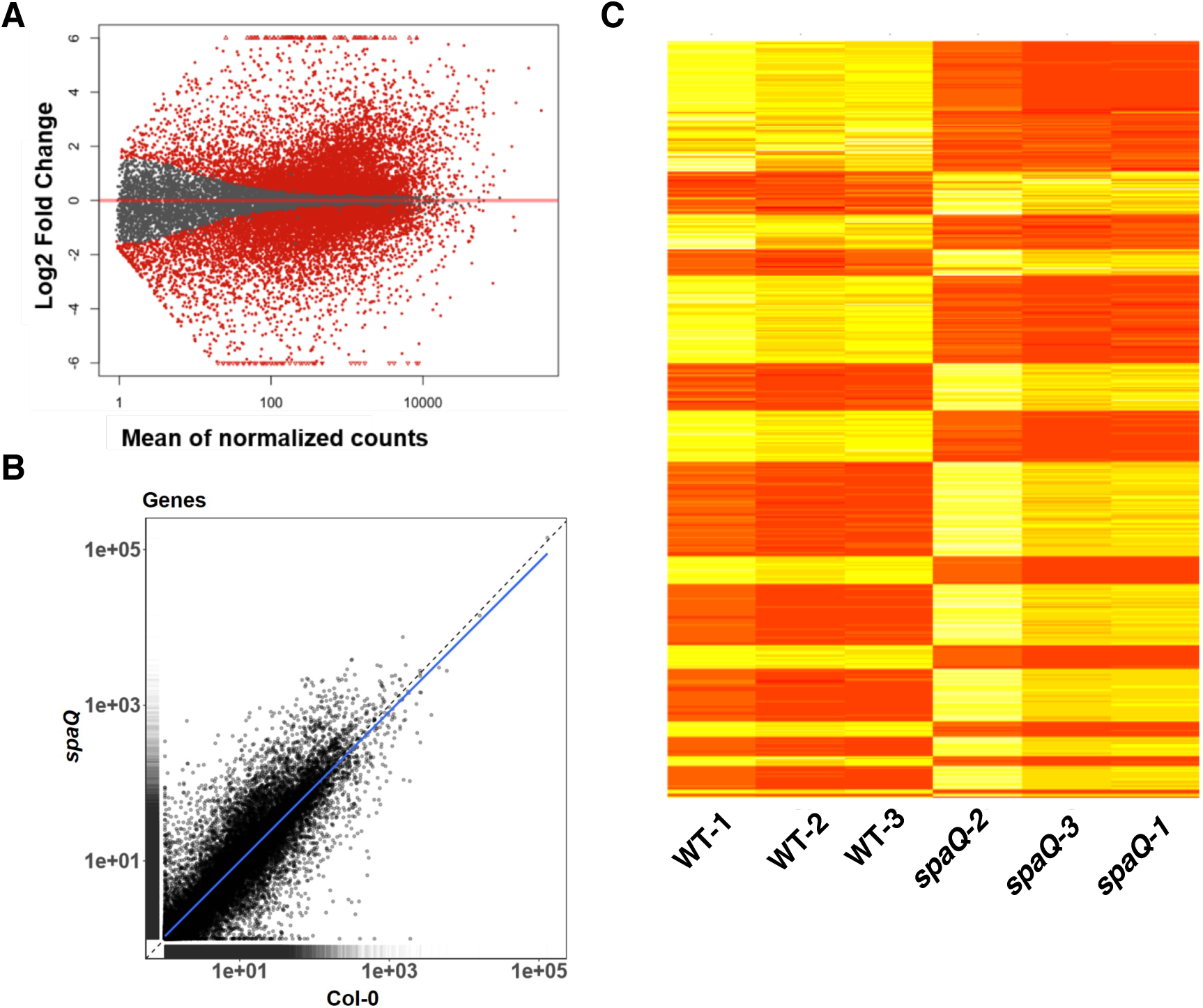
Genomic expression profile comparison between dark-grown wild-type (Col-0) and *spaQ* seedlings. (A) MA plot shows the log2 fold-change for the mean of normalized counts. The plot represents each gene with a dot. The x-axis presents the mean of normalized counts for all samples. The y-axis shows the log2 fold-change between *spaQ* mutant and wild-type in the dark. Genes with adjusted p-value below the threshold (alpha = 0.1) are shown in red. (B) A scatter plot shows the expression of genes in wild-type (Col-0) and *spaQ* mutant, with the x-axis representing the gene expression values for wild-type and the y-axis representing the gene expression values in *spaQ* mutant. Each point represents the expression of a gene in both backgrounds. Data generated from gene expression value from Cuffdiff analysis. The blue line indicates the linear regression line. (C) Heatmap represents the expression of 7261 differentially expressed genes in wild-type (WT-1,2,3) and *spaQ (spaQ-1,2,3*) mutants. Data generated from differential genes expressed in *spaQ* and raw counts from the three replicates of wild-type and *spaQ* mutants. Yellow indicates high expression and red indicates low expression.

The summaries of expression values of *spaQ* and Col-0 from three replicates for each gene are presented in the Scatter plot (Fig. 1B). Some common genes with similar expression values run along the diagonal line. However, we observed a very high number of genes differentially expressed in *spaQ* compared to wild-type. Using DESeq2, with the selected cut-off of log2 fold-change of 2 and FDR < 0.05, we identified 7261 differentially expressed genes (DEG) in *spaQ* mutant when compared to wild-type which were designated as *SPA-*regulated genes (gene listed with log2 fold-change values and adjusted p-values in Dataset S1). Hierarchical clustering of these 7261 genes is shown in Fig. 1C. Among 7261 genes, a large number of genes show distinct expression patterns in these two backgrounds. These data indicate that the *spaQ* mutations cause major changes in gene expression in dark-grown seedlings.

Biological functional category analysis revealed that most of genes involved in oxidation-reduction process, photosynthesis, defense response, response to cold, response to light stimulus and flavonoid biosynthetic process were up-regulated in *spaQ* mutants. Down-regulated genes are enriched in the terms related to response to auxin, abscisic acid, post-translational protein modification, regulation of transcription (Table 1; Fig. S5). The gene ontology analyses are well in line with the findings that SPAs function as a main regulator of seedling development, acting as an E3 ligase complex with COP1 to promote the ubiquitination and degradation of multiple substrates involved in seedling de-etiolation and photoperiodic flowering (Fittinghoff et al. 2006, Laubinger et al. 2006).

**Table 1.**
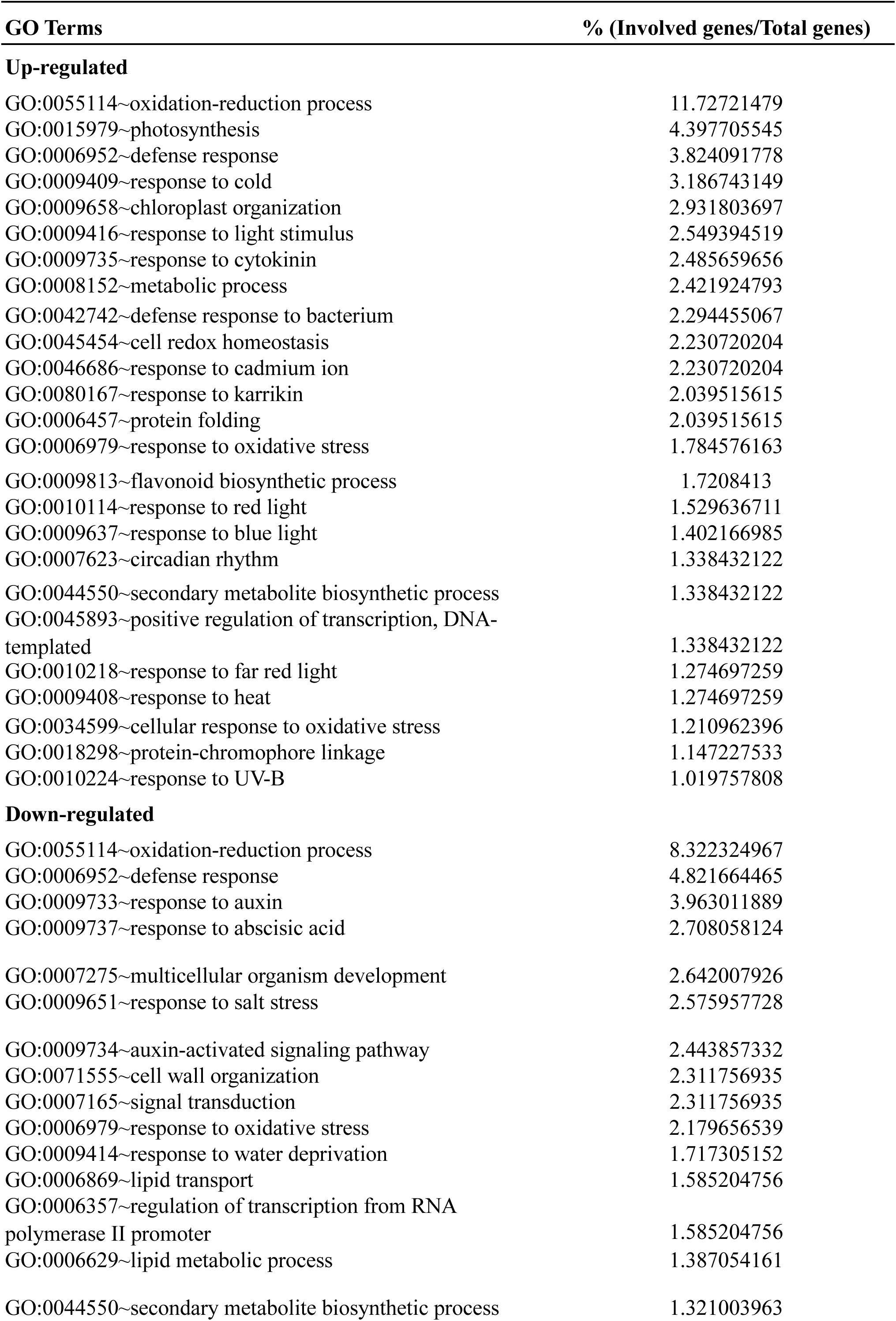
Summary of cellular and metabolic pathways regulated in *spaQ* mutant

| **GO Terms** | **% (Involved genes/Total genes)** |
| --- | --- |
| **Up-regulated** |  |
| GO:0055114~oxidation-reduction process | 11.72721479 |
| GO:0015979~photosynthesis | 4.397705545 |
| GO:0006952~defense response | 3.824091778 |
| GO:0009409~response to cold | 3.186743149 |
| GO:0009658~chloroplast organization | 2.931803697 |
| GO:0009416~response to light stimulus | 2.549394519 |
| GO:0009735~response to cytokinin | 2.485659656 |
| GO:0008152~metabolic process | 2.421924793 |
| GO:0042742~defense response to bacterium | 2.294455067 |
| GO:0045454~cell redox homeostasis | 2.230720204 |
| GO:0046686~response to cadmium ion | 2.230720204 |
| GO:0080167~response to karrikin | 2.039515615 |
| GO:0006457~protein folding | 2.039515615 |
| GO:0006979~response to oxidative stress | 1.784576163 |
| GO:0009813~flavonoid biosynthetic process | 1.7208413 |
| GO:0010114~response to red light | 1.529636711 |
| GO:0009637~response to blue light | 1.402166985 |
| GO:0007623~circadian rhythm | 1.338432122 |
| GO:0044550~secondary metabolite biosynthetic process | 1.338432122 |
| GO:0045893~positive regulation of transcription, DNA-templated | 1.338432122 |
| GO:0010218~response to far red light | 1.274697259 |
| GO:0009408~response to heat | 1.274697259 |
| GO:0034599~cellular response to oxidative stress | 1.210962396 |
| GO:0018298~protein-chromophore linkage | 1.147227533 |
| GO:0010224~response to UV-B | 1.019757808 |
| **Down-regulated** |  |
| GO:0055114~oxidation-reduction process | 8.322324967 |
| GO:0006952~defense response | 4.821664465 |
| GO:0009733~response to auxin | 3.963011889 |
| GO:0009737~response to abscisic acid | 2.708058124 |
| GO:0007275~multicellular organism development | 2.642007926 |
| GO:0009651~response to salt stress | 2.575957728 |
| GO:0009734~auxin-activated signaling pathway | 2.443857332 |
| GO:0071555~cell wall organization | 2.311756935 |
| GO:0007165~signal transduction | 2.311756935 |
| GO:0006979~response to oxidative stress | 2.179656539 |
| GO:0009414~response to water deprivation | 1.717305152 |
| GO:0006869~lipid transport | 1.585204756 |
| GO:0006357~regulation of transcription from RNA polymerase II promoter | 1.585204756 |
| GO:0006629~lipid metabolic process | 1.387054161 |
| GO:0044550~secondary metabolite biosynthetic process | 1.321003963 |

### SPA regulates the expression of a large group of transcription factors

SPA proteins have been shown to function as co-factors of the COP1 E3 ligase in controlling protein stability of a number of transcription factors (TF) in plants (Pham et al. 2018a). We compared the number of transcription factor genes significantly regulated by *spaQ* to the transcription factor database (http://planttfdb.cbi.pku.edu.cn/). Among a predicted total of 1717 transcription factor genes in Arabidopsis, there were 482 (∼28%) genes which exhibited more than 2-fold significant expression changes between *spaQ* and the wild-type. The description for the function of the transcription factors regulated by *spaQ* is also presented in Dataset S2.

We also identified 357 (∼21%) transcription factor genes differentially expressed in *cop1-4* when compared to the wild-type (see below), among which 300 overlapped between *cop1-4* and *spaQ* regulated transcription factors (Fig. S6; Dataset S2). However, 182 transcription factor genes differentially expressed in *spaQ* but not in *cop1-4.* A heat map shows the expression of the top 50 transcription factors that were differentially expressed in *spaQ* compared to WT (Fig. 2). The raw read count data for the transcription factors DEG in *spaQ* is shown in Dataset S2.

**Figure 2.**
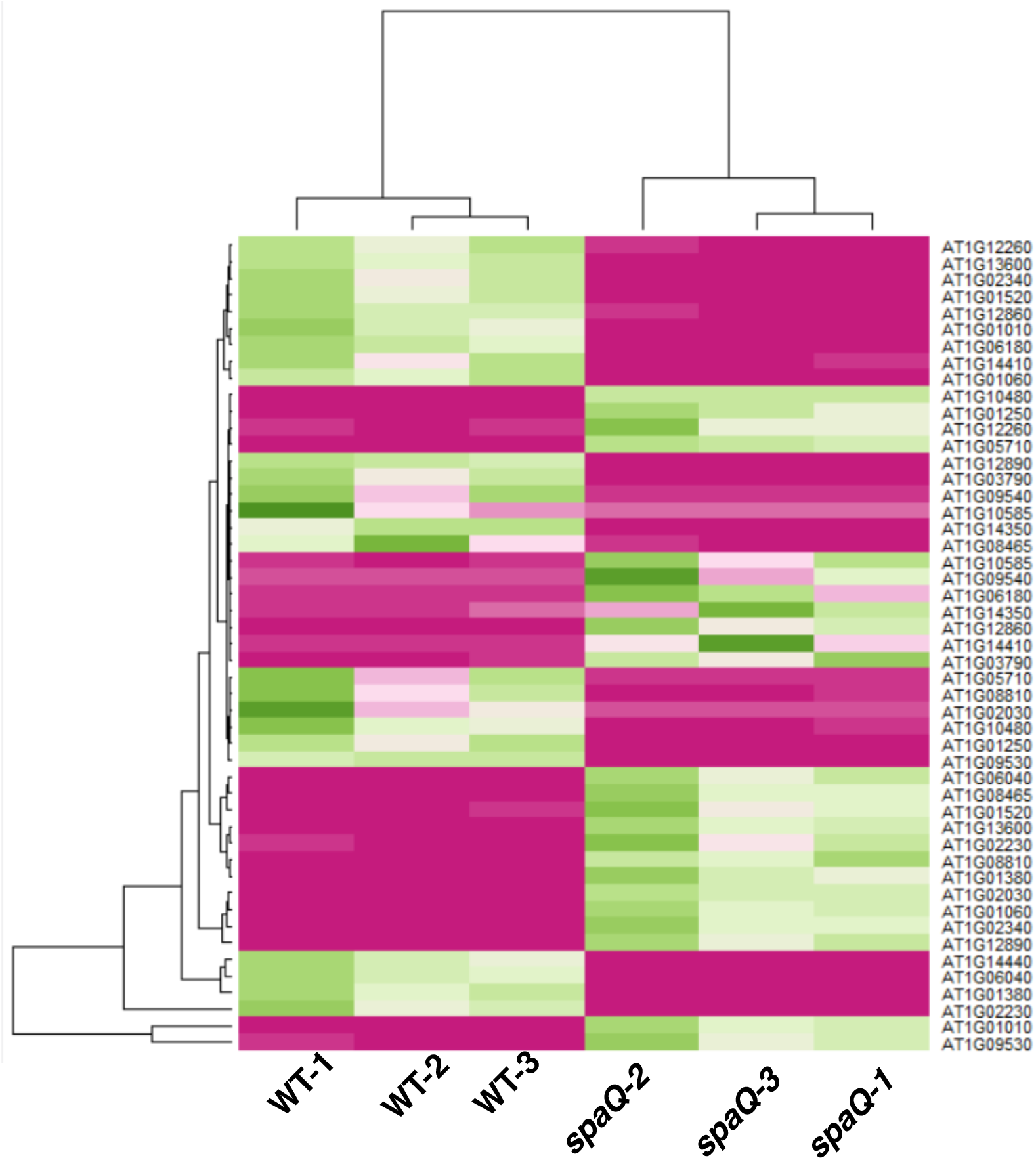
SPAs regulate the expression of a large group of transcription factor genes. Among 1717 transcription factor (TF) from the database, 482 TF were identified as significantly differentially expressed in *spaQ* mutant (Additional file 4). Color scale: pink: high expression, green: low expression. Heatmap represents the expression of top 50 TFs differentially expressed in *spaQ* compared to wild-type (WT) from three replicates, both up-regulated and down-regulated genes. RNA samples were extracted from 3-day-old dark-grown Col-0 and *spaQ* seedlings.

The significantly differential expression of transcription factors in *spaQ* suggests that SPA proteins play an important role in orchestrating the transcriptome by affecting the transcription factors. Gene Ontology (GO) analysis reveals that many regulated transcription factors involved in photomorphogenesis, cell-differentiation, ethylene activated response pathway, response to auxin, response to gibberellin, salt stress, ABA response and salicylic acid (Table 2). Many of the transcription factors belong to the bHLH and bZIP families that play central roles in photomorphogenesis. Interestingly, WUSCHEL-related homeobox genes were also up-regulated in *cop1-4* and *spaQ*. This is consistent with the previous study showing the function of light in activating the stem cell inducer *WUSCHEL* (Pfeiffer et al. 2016). In this case, the up-regulated expression of those WUSCHEL-related homeobox in *cop1-4* and *spaQ* mutants suggests a transition from dark to light-regulated development in *cop1-4* and *spaQ* even in the dark. Regulation of similar sets of transcription factors in *cop1-4* and *spaQ* suggests that these transcription factors might be the transcriptional targets of the COP1-SPA E3 ligase.

**Table 2.** Transcription factor genes regulated in *spaQ*

| **GO Term** | **% (Involved genes/Total genes)** |
| --- | --- |
| GO:0006355~regulation of transcription, DNA-templated | 85.06224066 |
| GO:0006357~regulation of transcription from RNA polymerase II promoter | 14.3153527 |
| GO:0030154~cell differentiation | 14.52282158 |
| GO:0009640~photomorphogenesis | 2.282157676 |
| GO:0009733~response to auxin | 6.22406639 |
| GO:0009873~ethylene-activated signaling pathway | 9.128630705 |
| GO:0009751~response to salicylic acid | 8.298755187 |
| GO:0009651~response to salt stress | 7.468879668 |
| GO:0009737~response to abscisic acid | 6.846473029 |
| GO:0009739~response to gibberellin | 6.43153527 |
| GO:0009753~response to jasmonic acid | 6.22406639 |
| GO:0009585~red, far-red light phototransduction | 1.867219917 |
| GO:0009723~response to ethylene | 4.771784232 |
| GO:0010200~response to chitin | 4.564315353 |
| GO:0045892~negative regulation of transcription, DNA-templated | 3.319502075 |
| GO:0009414~response to water deprivation | 3.319502075 |
| GO:0046686~response to cadmium ion | 3.319502075 |
| GO:0009740~gibberellic acid mediated signaling pathway | 2.697095436 |
| GO:0048366~leaf development | 2.697095436 |
| GO:0009908~flower development | 2.697095436 |
| GO:0000165~MAPK cascade | 2.489626556 |
| GO:0007623~circadian rhythm | 2.489626556 |
| GO:0009738~abscisic acid-activated signaling pathway | 2.489626556 |
| GO:2000652~regulation of secondary cell wall biogenesis | 2.282157676 |
| GO:0048510~regulation of timing of transition from vegetative to reproductive phase | 1.867219917 |
| GO:0048364~root development | 1.867219917 |
| GO:0010089~xylem development | 1.659751037 |
| GO:0009909~regulation of flower development | 1.659751037 |
| GO:0006970~response to osmotic stress | 1.659751037 |
| GO:0080167~response to karrikin | 1.659751037 |
| GO:0010187~negative regulation of seed germination | 1.452282158 |
| GO:0010218~response to far red light | 1.452282158 |
| GO:0009965~leaf morphogenesis | 1.452282158 |
| GO:0009736~cytokinin-activated signaling pathway | 1.452282158 |
| GO:0010228~vegetative to reproductive phase transition of meristem | 1.452282158 |
| GO:0009658~chloroplast organization | 1.452282158 |
| GO:0009641~shade avoidance | 1.244813278 |
| GO:0009741~response to brassinosteroid | 0.829875519 |
| GO:0048573~photoperiodism, flowering | 0.829875519 |
| GO:0009785~blue light signaling pathway | 0.622406639 |

### Genomic expression profile comparison between dark-grown *cop1-4* and *spaQ* seedlings

Genomic expression profiles regulated by various *cop1* alleles have been analyzed by RNAseq and microarrays (Ma et al. 2002, Wang et al. 2016). However, these studies were done using a condition which is not comparable to our study. To compare directly, we also sequenced *cop1-4* grown under identical conditions. By comparing the expression profiles of dark-grown *cop1-4* with wild-type (*cop1-4*/WT), we also revealed 5249 significantly DEGs with more than two-fold change and FDR < 0.05 in *cop1-4* (Dataset S1) which are designated as *cop1-4* regulated genes. We examined the overlap of the differentially expressed genes between *cop1-4* and *spaQ* mutants. Venn diagrams show that among 5249 gene differentially expressed in *cop1-4*, 4471 (85.17%) genes were co-regulated also in *spaQ* (Dataset S3, Fig. 3A). Among these 4471 co-regulated genes, 2318 genes were co-upregulated and 2137 genes were co-downregulated. Furthermore, hierarchical clustering reveals that these genes are co-regulated in the same direction in *cop1-4* and *spaQ* mutants (Fig. 3B). Indeed, the scatterplot shows very strong co-expression profile between *cop1-4* and *spaQ* mutants (the Pearson correlation coefficient *r*= 0.95) (Fig. 3E). We also performed gene ontology (GO) enrichment analysis to identify biological processes (Huang et al. 2009), in which the genes co-regulated between *cop1-4* and *spaQ* are involved. GO analysis reveals that the differentially expressed genes (DEGs) can be classified into 172 GO terms (Fig. 3C, D, Dataset S3). The up-regulated genes are classified into cell redox homeostasis, chlorophyll biosynthetic process, photosynthesis regulated genes, response to light stimulus (majority in red light and small fraction of far-red light responsive genes). Especially, 66 genes (∼3%) involved in response to cold and embryo development resulting in seed dormancy. This is consistent with the previous study showing the function of COP/SPA as E3 ligase for HY5 and negatively regulate the gene expression of cold-responsive genes as previously shown (Catalá et al. 2011).

The down-regulated genes in *cop1-4* and *spaQ* include cell wall organization, regulation of growth, regulation of transcription, response to auxin, and response to abscisic acid. Interestingly, 4.5% of total down-regulated genes belong to leucine-rich receptor-like protein kinase family protein, regulation of transcription, and other hormone, especially abscisic acid, gibberellic acid, and response to stress.

**Figure 3.**
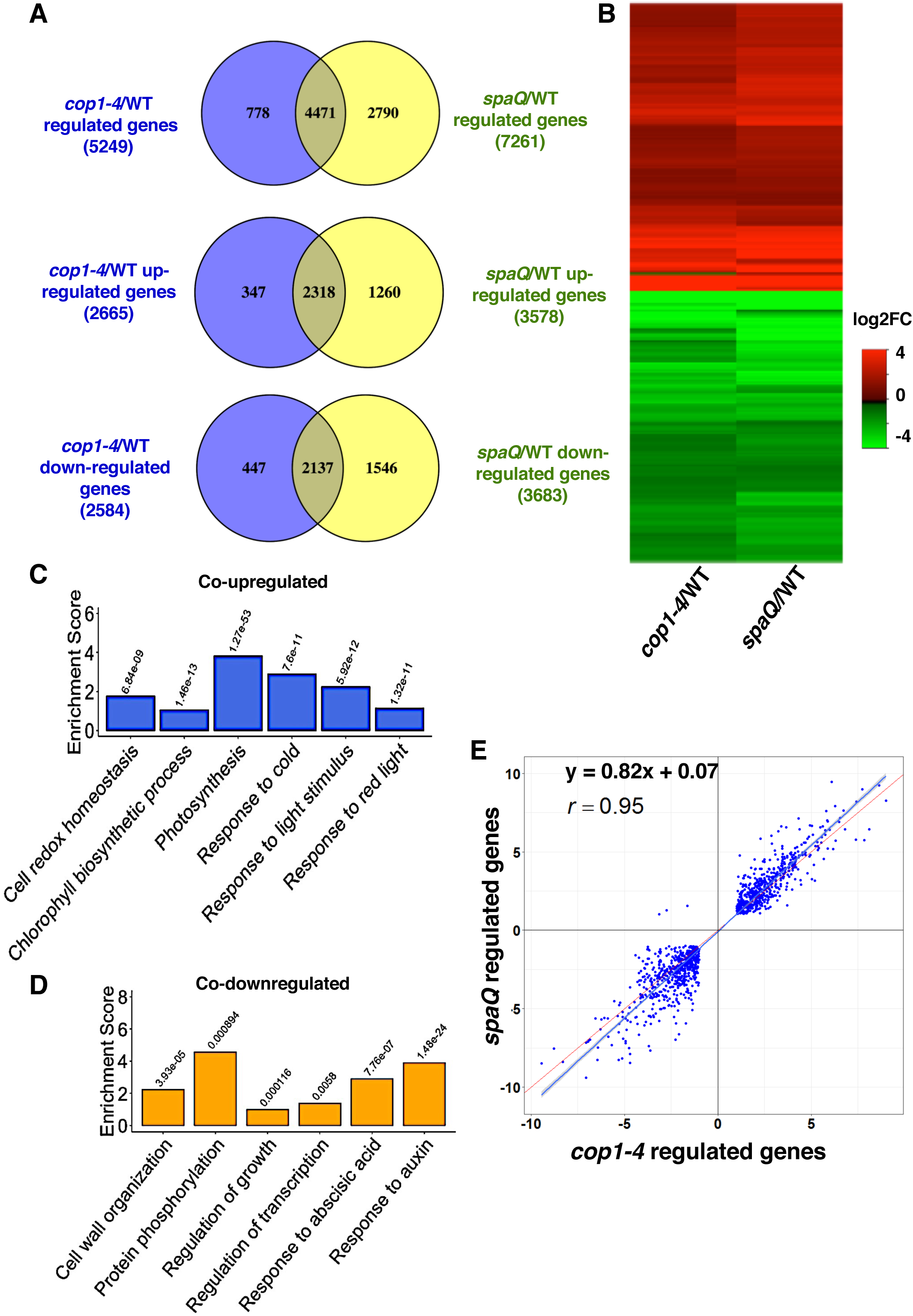
Genomic expression profile comparison between 3-day old dark-grown *cop1-4* and *spaQ* seedlings. (A) (B) Venn diagram (A) and Hierarchical clustering (B) of 4471 differentially expressed genes (DEG) in two different pairwise comparisons indicated (*cop1-4*/WT and *spaQ*/WT). Among 4471 DEGs, 2318 genes are co-upregulated and 2137 genes are co-down-regulated (A). Genes showing differential expression in both pairwise comparisons are presented (fold change >2, FDR < 0.05). Number of genes up-regulated and down-regulated are shown. Color scale displays log2 fold-change (log2FC) value of significantly differential expression. (C) to (D) Bar graphs showing GO enrichment analysis of genes which are significantly co-upregulated (C), and co-downregulated in *cop1-4/*WT and *spaQ*/WT. Enrichment scores represent the percentage (Involved genes/Total genes). p-value was shown for each GO term. (E) Pearson correlation scatter plot (*r* = 0.95, p-value < 2.2e-16) shows a high correlation between *cop1-4* and *spaQ* regulated genes.

Among 4471 genes co-regulated by *cop1-4* and *spaQ*, we also found 1396 genes (31%) to share the similar expression profile with light-grown wild-type seedlings (Fig. S7). Expression profiles of light-grown wild-type seedlings were obtained from a previous study (Dong et al. 2014), in which the seedlings were grown for 4 days in the dark and then expose for 6hrs of white light.

Previous study has shown that DET1 interacts with the key transcription factors and regulates transcription of thousands of downstream genes in Arabidopsis to repress photomorphogenesis (Dong et al. 2014). Therefore, we also compared genomic profiles of our data for *cop1-4* and *spaQ* with *det1-1* mutant profiles. *det1-1* mutant was grown in dark for 4 days and then exposed to white light for 6h. Using the DEGs in *det1-1* mutant, we found that 1967 genes are significantly co-regulated by *cop1-4* and *spaQ* (Fig. S8). Moreover, the hierarchical clustering illustrates highly similar patterns between these three mutants. These data suggest a shared function of COP1, SPA and DET1 as central repressors of light signaling that regulate large changes in gene expression in the dark to promote skotomorphogenesis.

### Comparison of differential genes between *cop1-4*, *spaQ* and *hy5* mutants

HY5 is well-known as one of the direct targets of the COP1-SPA E3 ligase in the dark involved in light-mediated transcriptional regulation. COP1-SPA promotes HY5 degradation in the dark, thereby, inhibiting light-activated gene expression in darkness. To examine the relationships among these three components, we compared the gene expression profiles among these three mutants. We extracted the raw count from RNA-Seq data obtained from the *hy5-215* mutant grown in the light (Zhang et al. 2011). *hy5-215* seeds were incubated at 4°C in the dark for 4 days, after which they were exposed to continuous white light (WL, 170 µmol m^−2^ sec^−1^) at 22°C for 6 days. From the raw count, we analyzed the differential gene expression in the *hy5* mutant compared to wild-type under the same condition previously described using DESeq2 (Zhang et al. 2011), called *hy5*-regulated genes *(hy5*/WT) (Dataset S4). Indeed, comparing *cop1-4* and *spaQ* genomic expression profiles with the gene expression profile in *hy5* mutant, 1060 overlapping genes are expressed in antagonistic manner, as shown in hierarchical clustering (Fig. 4A, D).

**Figure 4.**
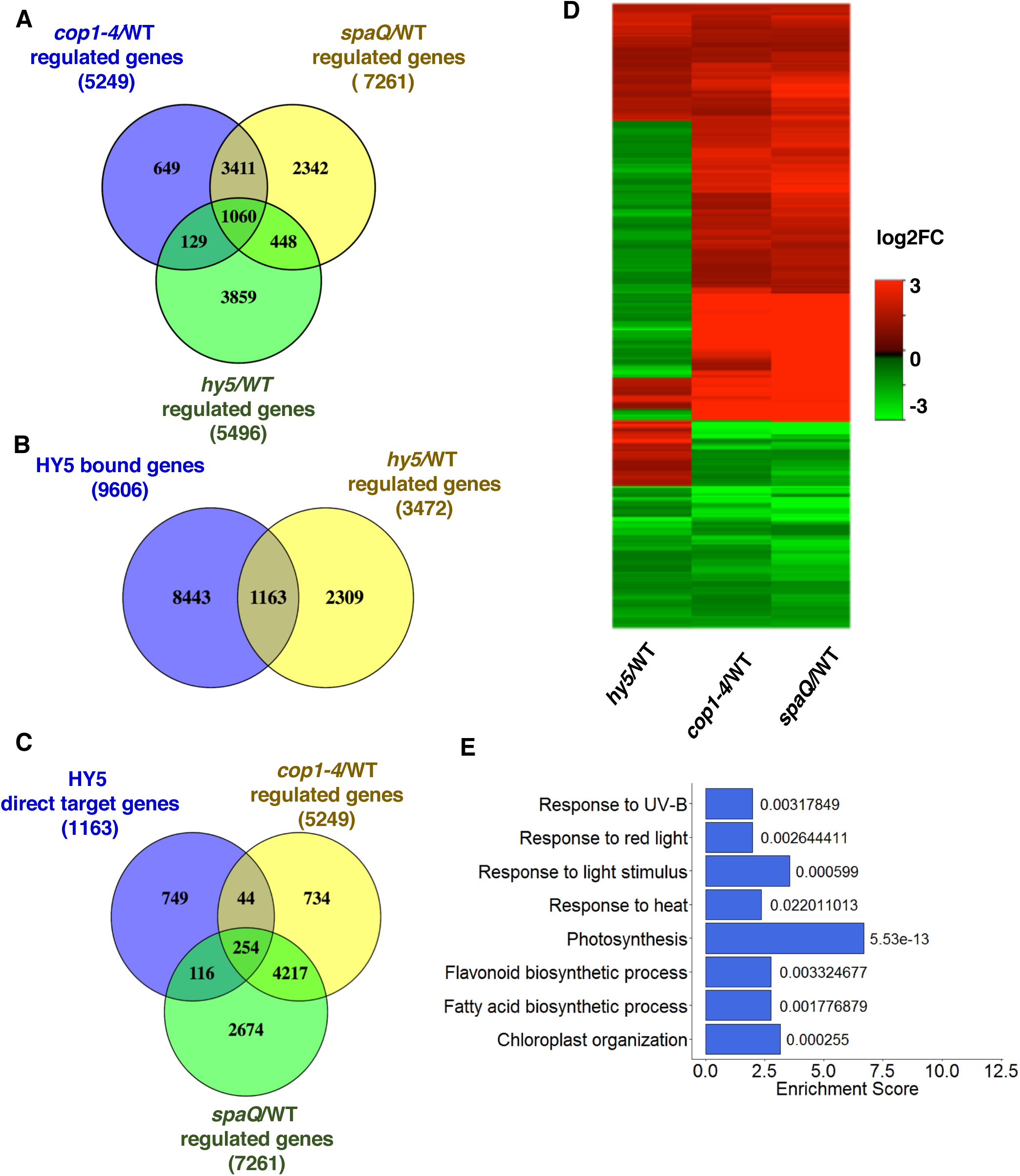
Comparison of gene expression profiles between *cop1-4*, *spaQ* and *hy5* mutants. (A) Venn diagram shows the comparison of 5496 regulated genes in *hy5* with those in *cop1-4* and *spaQ*. 1060 genes co-regulated by *cop1-4, spaQ* and *hy5* mutants are presented. Col-0 *cop1-4* and *spaQ* seedlings were grown in 3 days in dark. *hy5* mutant was grown 4 days under white light. (B) Venn diagram shows the comparison of 9606 HY5 bound genes and 3472 *hy5*/WT significantly regulated genes (Zhang et al. 2011). 1163 overlapped genes are referred as HY5 direct target genes. (C) Venn diagram shows the comparison of 1163 HY5 direct target genes and *cop1-4*/WT and *spaQ*/WT regulated genes. (D) Heatmap shows clustering expression patterns of 254 HY5 direct target genes that are co-regulated by *cop1-4* and *spaQ*. (E) Bar graphs show GO enrichment analysis of genes which are HY5 direct target genes and significantly co-regulated in *cop1-4/*WT and *spaQ*/WT. Enrichment scores represent the percentage (Involved genes/Total genes). p-value is shown for each GO term.

From 5496 HY5-regulated genes in Figure 4A, we selected 3472 genes regulated by HY5 which are differentially expressed more than 2-fold in *hy5-215* mutant (Dataset S5). We also compared the 9606 HY5-bound genes under white-light (Zhang et al. 2011) with those of the 3472 *hy5*-regulated genes to obtain 1163 overlapped genes that are referred as HY5-direct target genes (Fig. 4B). By comparing the 1163 HY5-direct target genes with the *cop1-4*- and *spaQ*-regulated genes, we identified 254 overlapping genes which are co-regulated in *cop1-4, spaQ* and *hy5* (Fig. 4C) (Dataset S6). In fact, a large number of genes differentially expressed more than 2-fold in *cop1-4* and *spaQ,* but expressed less than two-fold in *hy5* mutant. These data are also consistent with previous *cop1* microarray data (Ma et al. 2002). GO analysis shows that those genes largely belong to photosynthesis, flavonoid biosynthetic, chloroplast organization and response to light stimulus (Fig. 4E).

Interestingly, we also compared genes differentially expressed in *det1-1* mutant with the *hy5* regulated genes and found antagonistic expression of *hy5* regulated genes in *det1-1* mutant similar to *cop1* and *spaQ* (Fig. S9). Thus, these three constitutively photomorphogenic mutants display opposite expression patterns compared to *hy5*, as established by previous biochemical and genetic analysis.

To complete the comparison with the HY5-direct target genes, we performed qRT-PCR to confirm the expression of HY5-direct target genes including HY5-upregulated genes (*ELIP2, CAB3, CHS* and *RBCS1A*) and HY5-down regulated genes (*PER59* and *PRP2*). qRT-PCR data largely reproduced the RNA-Seq data (Fig. 5). Since SPA proteins function as negative regulator of HY5 protein stability in the dark, gene expression analysis of HY5-target genes in *spaQ* showed significant rescue compared to wild-type and *hy5* mutant, except in the case of *PER59*. Overall, these data show that our RNAseq data can be verified by independent methods.

**Figure 5.**
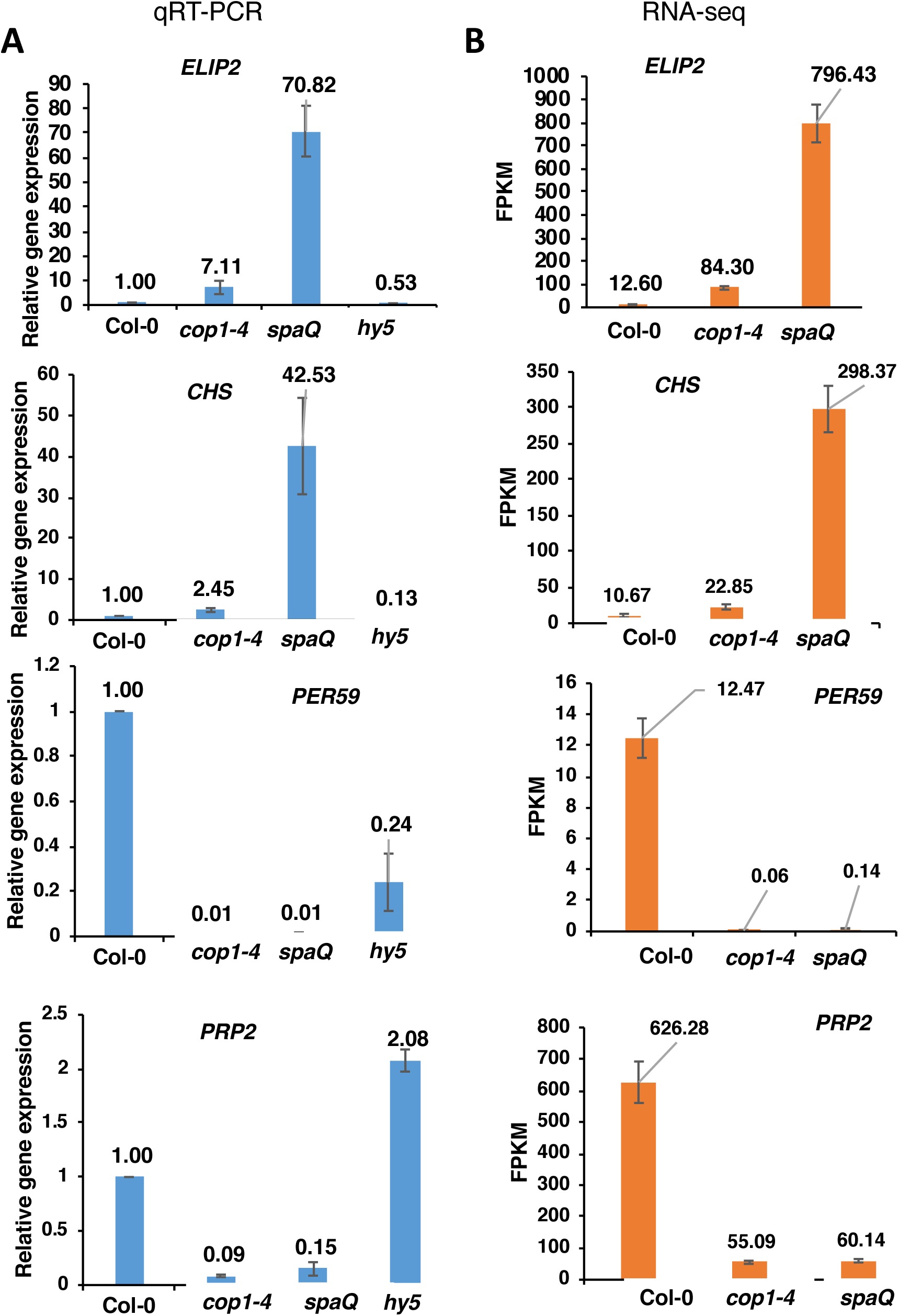
Expression levels of HY5 target genes in wild-type (Col-0), *cop1-4*, *spaQ*, and *hy5-215* mutants. (A) Expression level data using qRT-PCR. RNA was extracted from 3-day dark-grown seedlings of the indicated mutants. The relative gene expression level was calculated based on the *PP2A* gene expression. The expression level of wild-type (Col-0) was set as 1. (B) The normalized expression level (FPKM; fragment per kilobase per million reads) from RNA-Seq data.

### SPAs may function in COP1-dependent and -independent manners

It is unknown whether SPA proteins can function only in association with COP1 and/or they have a COP1-independent role in regulating plant development. Based on our differential gene expression analysis, we have examined whether some of the genes are prominently regulated only in *spaQ* or in *cop1-4*. We have identified a list of genes and GO terms that are predominantly regulated by SPAs or COP1 including transcription factor genes (Dataset S7-S8 and Figs. S10-S11). Enrichment of distinct GO terms in *spaQ-* and *cop1*-regulated genes suggests that these factors might have independent roles in regulating plant responses and developmental pathways.

A closer scrutiny revealed that most of the genes differentially expressed in *spaQ* mutant are involved in response to heat, embryo development, oxidation–reduction process, defense response, and protein phosphorylation (Fig. S10). Interestingly, we identified a master regulator of heat responsive genes which is significantly down-regulated in *spaQ*. For example, *GASA5* was shown as the negative regulator in thermotolerance in Arabidopsis by regulating both salicylic acid (SA) signaling and heat shock-protein accumulation (Zhang et al. 2011). As a consequence, a number of heat shock genes (*HSP90.1* and *HSP70*) are up-regulated in *spaQ* mutants. However, those genes are not significantly regulated in *cop1-4* and *cop1-6* alleles (Fig. 6). We also found that the cold and ABA inducible gene, *KIN1*, is strongly up-regulated in the *spaQ* mutant, but not significantly in *cop1-4* and *cop1-6* mutants. The expression levels of *KIN1*, *GASA5* and heat shock genes in *spaQ* and different *cop1* mutants (*cop1-4* and *cop1-6*) were verified by qRT-PCR (Fig. 6). Overall, these data suggest that SPAs may have COP1-independent roles in regulating plant responses to various cues. However, further analyses are necessary to support this conclusion.

**Figure 6.**
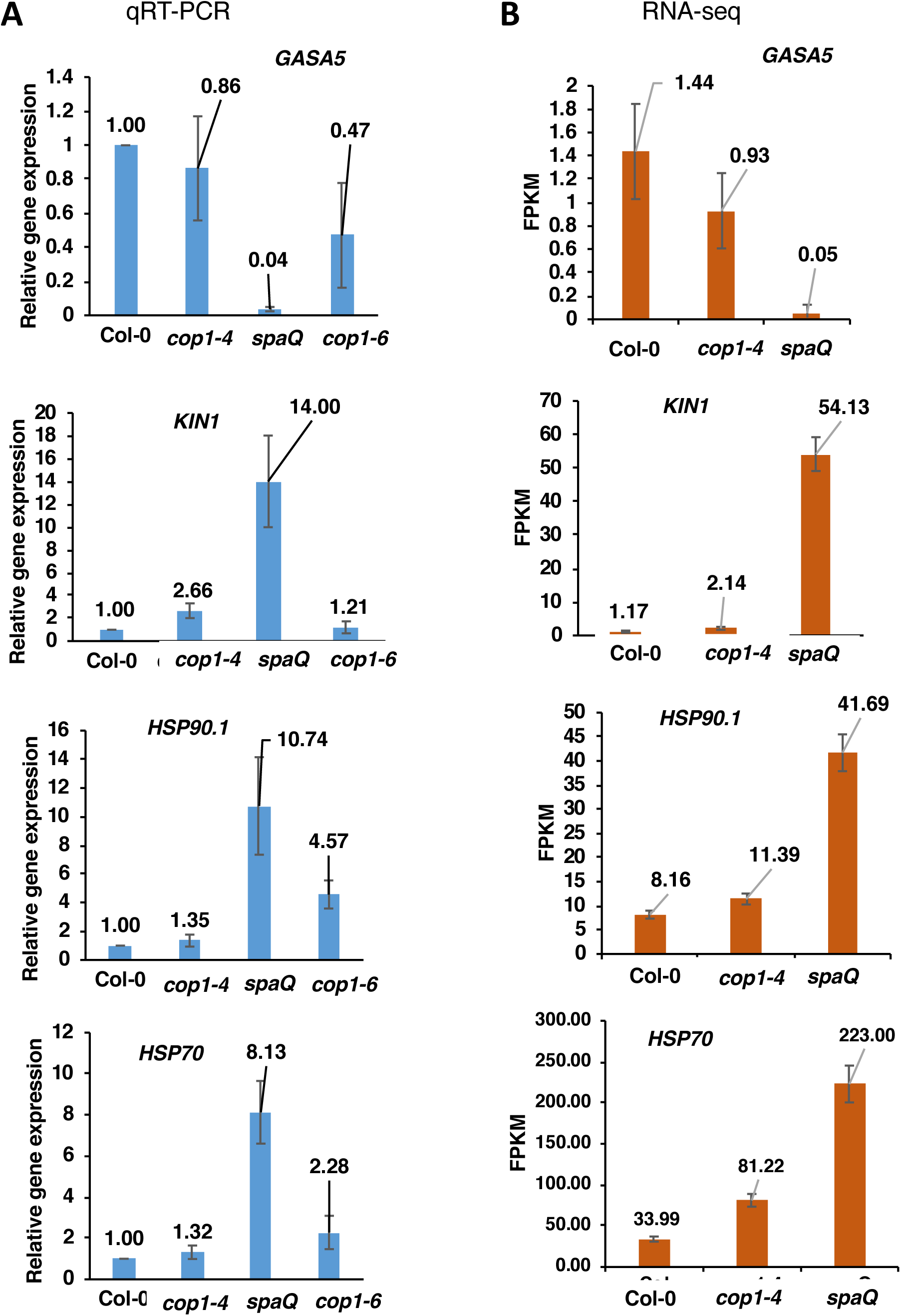
Expression level of some genes differential expressed in *spaQ* mutant, compared to wild-type, *cop1-4* and *cop1-6* mutants. (A) expression level using qRT-PCR. RNA was extracted from 3-day dark-grown seedlings. The relative gene expression level was calculated based on the *PP2A* gene expression. Expression level of wild-type (Col-0) was set as 1. (B) The normalized expression level (FPKM; fragments per kilobase per million reads) from RNA-Seq data.

Interestingly, we also identified a few genes that are down-regulated predominantly in *cop1-4,* but not in *spaQ* (Dataset S7-S8; Figs. S12). These genes belong to the ABA inducible genes accumulated during seed maturation (AT5G44120, AT4G28520) and late embryogenesis abundant proteins (LEA) (AT3G51810, EM1; AT2G40170, EM6; AT3G17520, AT3G15670, AT2G42560, AT1G52690, AT5G44310) (see more in Dataset S5) (Hundertmark and Hincha 2008). However, some members of the same gene family (AT2G42540, AT2G03850, AT1G64065, AT5G22870) also display significantly different expression in *spaQ* mutant (Dataset S7-S8). These data suggest that both COP1 and SPA proteins regulate seed maturation, but they do so by regulating distinct members of the same gene family.

## Discussion

Previously, extensive genetic, biochemical and molecular analyses revealed the involvement of SPA proteins in regulating photomorphogenesis and other biological processes (Hoecker 2017). In this study, we show the genomic bases for the SPA-regulated plant developmental pathways. Strikingly, more than 26% of the Arabidopsis genes are regulated by SPA proteins in 3-day-old dark-grown seedlings. This is close to the ∼30% of the Arabidopsis genes regulated by light signals (Ma et al. 2001). Because SPA proteins mostly function in the dark, this mirror image pattern of gene expression highlights the crucial roles played by SPA proteins in regulating photomorphogenesis.

GO term analyses revealed that photosynthesis, chloroplast organization and responses to red, far-red and blue lights are upregulated, while cell wall organization, and response to auxin are downregulated in the *spaQ* mutant compared to wild-type. This is consistent with the previous genetic and biochemical analyses showing that SPA proteins repress photomorphogenesis and promote cell elongation in darkness (Laubinger et al. 2004).

Comparison of the gene expression patterns in *cop1-4*, *det1* and *spaQ* mutants reveal a striking similarity among all three constitutively photomorphogenic mutants. Conversely, gene expression patterns in the *hy5* mutant are antagonistic to those in the *cop1-4*, *det1* and *spaQ* mutants. Because HY5 is a positively acting transcription factor promoting photomorphogenesis, these data suggest that the negative regulators largely function by controlling the abundance and/or activity of the positive regulators like HY5. Moreover, only ∼20% of the *spaQ*-regulated genes overlap with *hy5*-regulated genes, suggesting that SPA proteins control gene expression not only through HY5, but also through other transcription factors. These data reflect the growing list of transcription factors that are regulated by COP1-SPA complex in darkness (Xu et al. 2015, Hoecker 2017).

Previous genetic and biochemical data revealed that SPA proteins mostly function in association with COP1, where *SPA1* and *SPA2* have a predominant function at the seedling stage, while *SPA3* and *SPA4* function in particular in adult plants (Hoecker 2017). The four SPA proteins interact with COP1 to form distinct COP1-SPA complexes in a manner that primarily depends on the respective abundance of the SPA proteins in different tissues and at different developmental stages (Zhu et al. 2008). Phenotypic analyses of *cop1* and *spa* mutants indicate a strong overlap of COP1 and SPA functions in the regulation of skotomorphogenesis, shade avoidance response and flowering time (Hoecker 2017). *cop1-5 spaQn* quintuple null mutants visually resemble *cop1-5* null mutants, suggesting that SPA proteins have no activity in the absence of COP1 (Ordoñez-Herrera et al. 2015). Our genomic data provide the evidence that these proteins might have overlapping as well as independent roles. Firstly, SPA proteins regulate significantly larger number of genes (7261 genes) compared to COP1 (5249 genes). In addition, while 85% of the *cop1-4*-regulated genes overlap with *spaQ*-regulated genes with very high correlation coefficient (r=0.95), only >62% of the *spaQ*-regulated genes overlap with *cop1*-regulated genes. In total this suggests that SPA proteins may regulate gene expression also independently of COP1. However, we cannot exclude the possibility that the use of weak alleles of *cop1* which retain partial COP1 activity (McNellis et al. 1994) caused the failure to observe a 100% overlap between *COP1*- and *SPA*-regulated genes. In particular, in contrast to our results, it was described previously that *CHS* and *CHI* genes are misregulated in *cop1* mutants (Deng et al. 1991, Ma et al. 2002). Null alleles of *cop1* (e.g. *cop1-5*) cause extremely severe phenotypes at the seedling stage including seedling arrest, thus restricting the use of such alleles for transcriptome experiments. It is possible that most of the *cop1*-independent genes in *spaQ* set would also be regulated by *COP1* if a *cop1* null allele had been used in these experiments. Nevertheless, GO analyses revealed COP1-independent roles of SPA proteins and SPA-independent roles of COP1 for specific ontologies. For example, *spaQ*-regulated genes are enriched with GO terms such as “response to heat”. Consistently, heat shock genes are upregulated in *spaQ*. Similarly, seed maturation and late embryogenesis abundant protein genes (*LEA*) are down-regulated in *cop1-4*. However, other members of the same family are regulated by SPA proteins. Although, these gene expression analyses suggest that COP1 and SPA proteins may have independent roles in regulating plant development, further phenotypic analyses are necessary to draw robust conclusion.

In summary, the transcriptome analyses presented here provide the molecular bases for the SPA-regulated developmental pathways in Arabidopsis. Most DEGs are co-regulated by COP1 and SPA proteins, thus confirming the actions of these proteins in a COP1/SPA complex. Further analysis of *cop1* null mutants as well as *spa* single, double, triple and quadruple mutants at different developmental stages might answer these questions.

## Author contributions

V.N.P., I.P., U.H. and E.H. designed experiments. V.N.P. and I.P. carried out experiments. V.N.P., I.P., U.H. and E.H. analyzed data and interpreted the results. V.N.P. wrote the article. V.N.P., I.P., U.H. and E.H. edited the manuscript.

## Acknowledgments

We thank Dr. Xing Wang Deng for sharing *cop1* mutant, and the Huq lab members for the technical support and critical reading of the manuscript. The authors acknowledge the Texas Advanced Computing Center (TACC) at The University of Texas at Austin for providing high performance computing, visualization, and database resources that have contributed to the research results reported in this paper. This work was supported by grants from the National Institute of Health (NIH) (GM-114297) and National Science Foundation (MCB-1543813) to E.H.

## Availability of data and materials

The data that support the findings of this study are openly available in Gene Expression Omnibus database at https://www.ncbi.nlm.nih.gov/geo/query/acc.cgi?acc=GSE112662, reference number GSE112662.

## Supporting information

**Table S1.** RNA-seq reads and mapping statistics.

**Table S2.** List of primers using for qRT-PCR.

**Figure S1.** Heat map of the sample-to-sample distances.

**Figure S2**. Principle component analysis (PCA).

**Figure S3.** Matrix of scatter plots for visualization of gene expression in wild-type (Col), *cop1-4* and *spaQ* mutants using Cufflinks and CummeRbund.

**Figure S4.** Volcano plots for visualizing significantly differential expressed genes in wild-type (Col), *cop1-4* and *spaQ* mutants using Cufflinks and CummeRbund.

**Figure S5.** Functional classification of SPA-regulated genes.

**Figure S6.** Venn diagram shows the comparison between Transcription factor (TF) regulated in *cop1-4* and *spaQ* mutants.

**Figure S7.** Comparison of gene expression profiles between light-grown wild-type seedlings, *cop1-4, and spaQ* in the dark.

**Figure S8.** Genomic expression profile comparison between dark-grown *cop1-4*, *spaQ,* and *det1-1* seedlings.

**Figure S9.** Comparison of gene expression profiles between *cop1-4, spaQ, det1-1,* and *hy5* mutants.

**Figure S10.** GO analysis of 2790 genes which are differentially expressed in *spaQ* mutant but not in *cop1-4* mutant.

**Figure S11.** GO analysis of 778 genes which are differentially expressed only in *cop1-4* mutant but not in *spaQ*.

**Figure S12.** The normalized expression level (FPKM; fragments per kilobase per million reads) of some differential expressed genes in *cop1-4* mutant, compared to wild-type (Col-0) and *spaQ* from RNA-Seq data.

**Dataset S1:** List of gene regulated by *cop1-4* and *spaQ* in the dark.

**Dataset S2:** List of transcription factor differential expressed in *spaQ*, *cop1-4* and GO terms analysis for TF regulated by *spaQ*.

**Dataset S3:** List of genes co-regulated by *cop1-4* and *spaQ* in the dark. Dataset S4: List of gene regulated by *hy5*

**Dataset S5:** List of genes significantly regulated by *hy5*

**Dataset S6:** List of 254 overlapping genes which are co-regulated in *cop1-4, spaQ* and *hy5*

**Dataset S7:** List of genes differential expressed in *spaQ* but not in *cop1-4* and vice versa.

**Dataset S8:** GO Terms analysis of gene differential expressed in *spaQ* but not in *cop1-4* and vice versa.

