## Supplemental figures for "Genomic evidence reveals SPA-regulated developmental and metabolic pathways in dark-grown *Arabidopsis* seedlings"

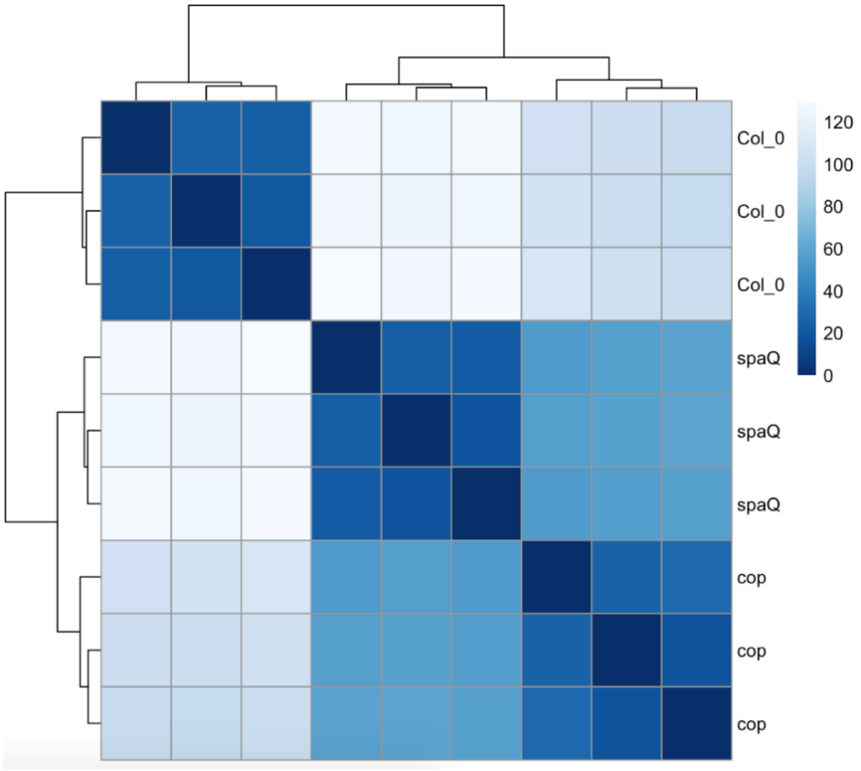


**Figure S1. Heat map of the sample-to-sample distances. Col-0: wild-type, *spaQ: spaQ* mutants, cop: *cop1-4* mutant**

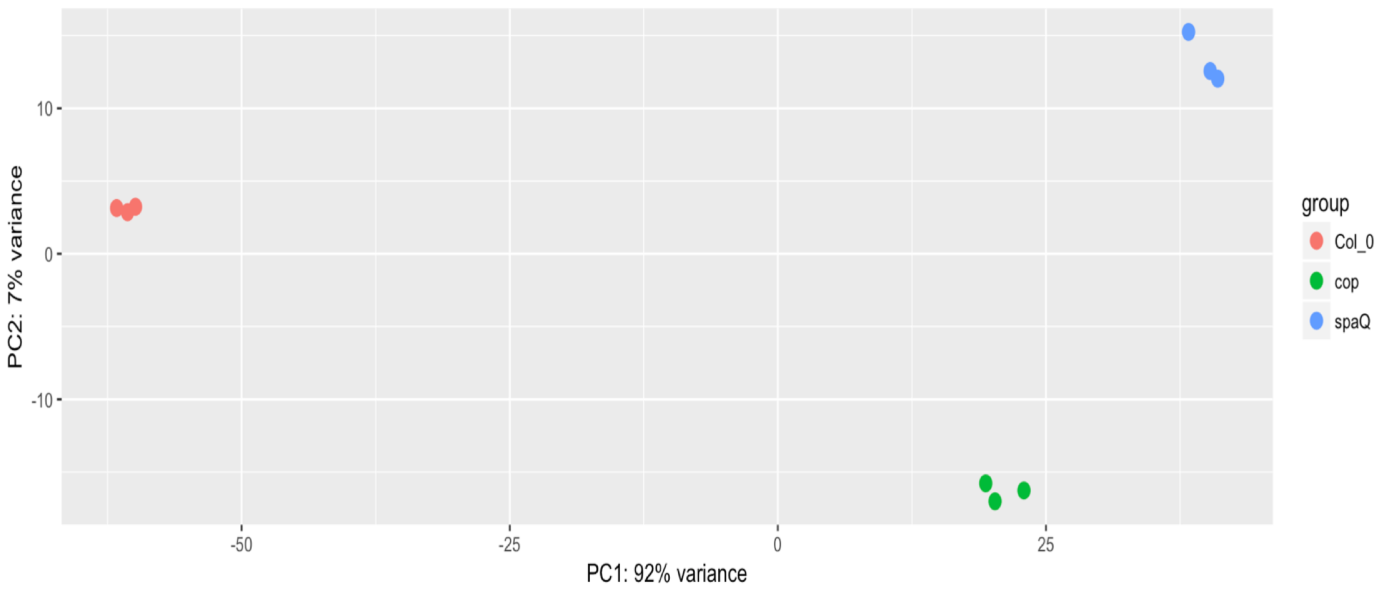


**Figure S2. Principle component analysis (PCA). Col-0: wild-type, *spaQ: spaQ* mutants, cop: *cop1-4* mutant**

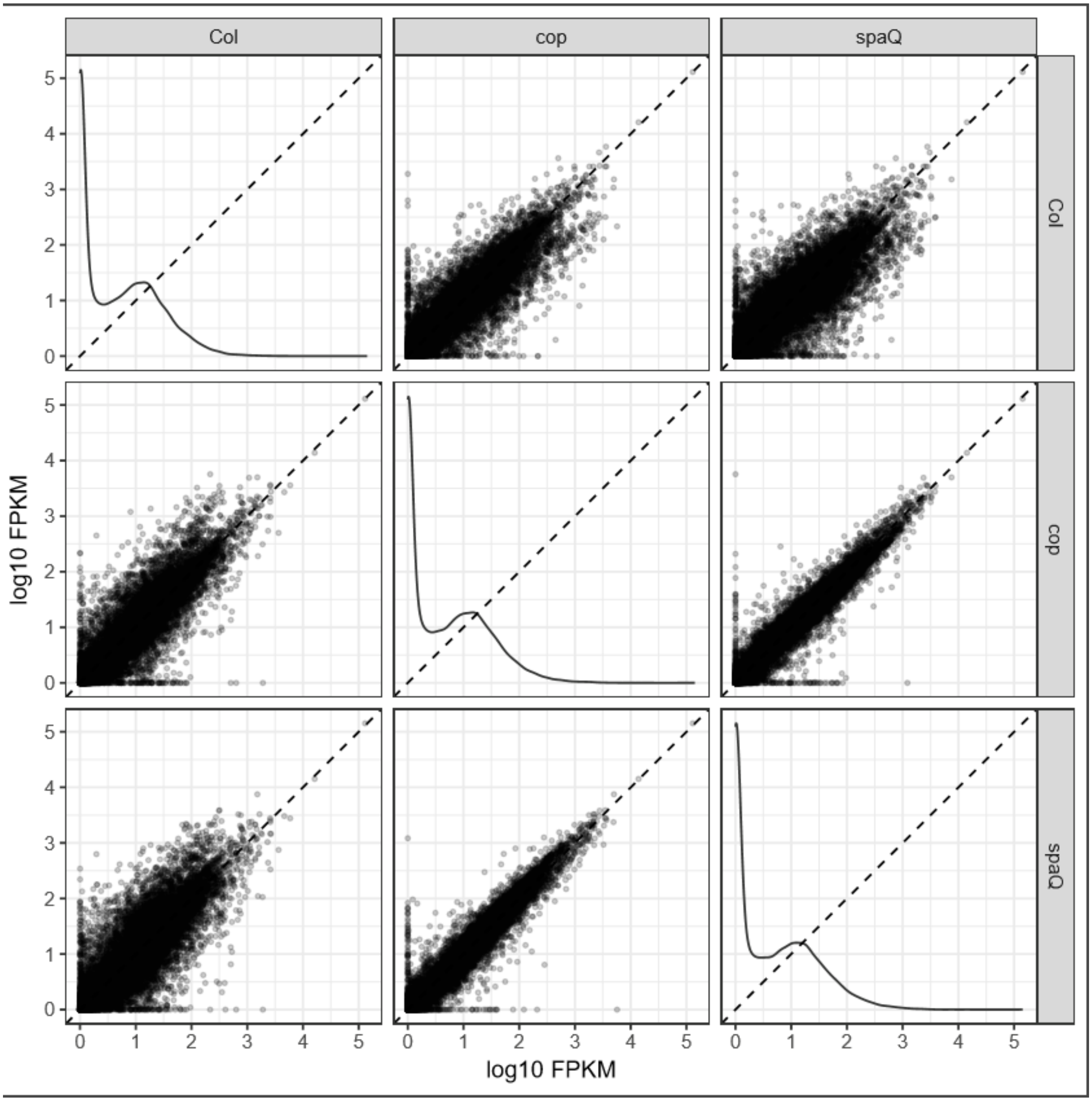


**Figure S3. Matrix of scatter plots for visualization of gene expression in wild type (Col), *cop1-4* and *spaQ* mutants using Cufflinks and CummeRbund.** The normalized expression level (FPKM; Fragment per kilobase per million reads) from RNA-seq data.

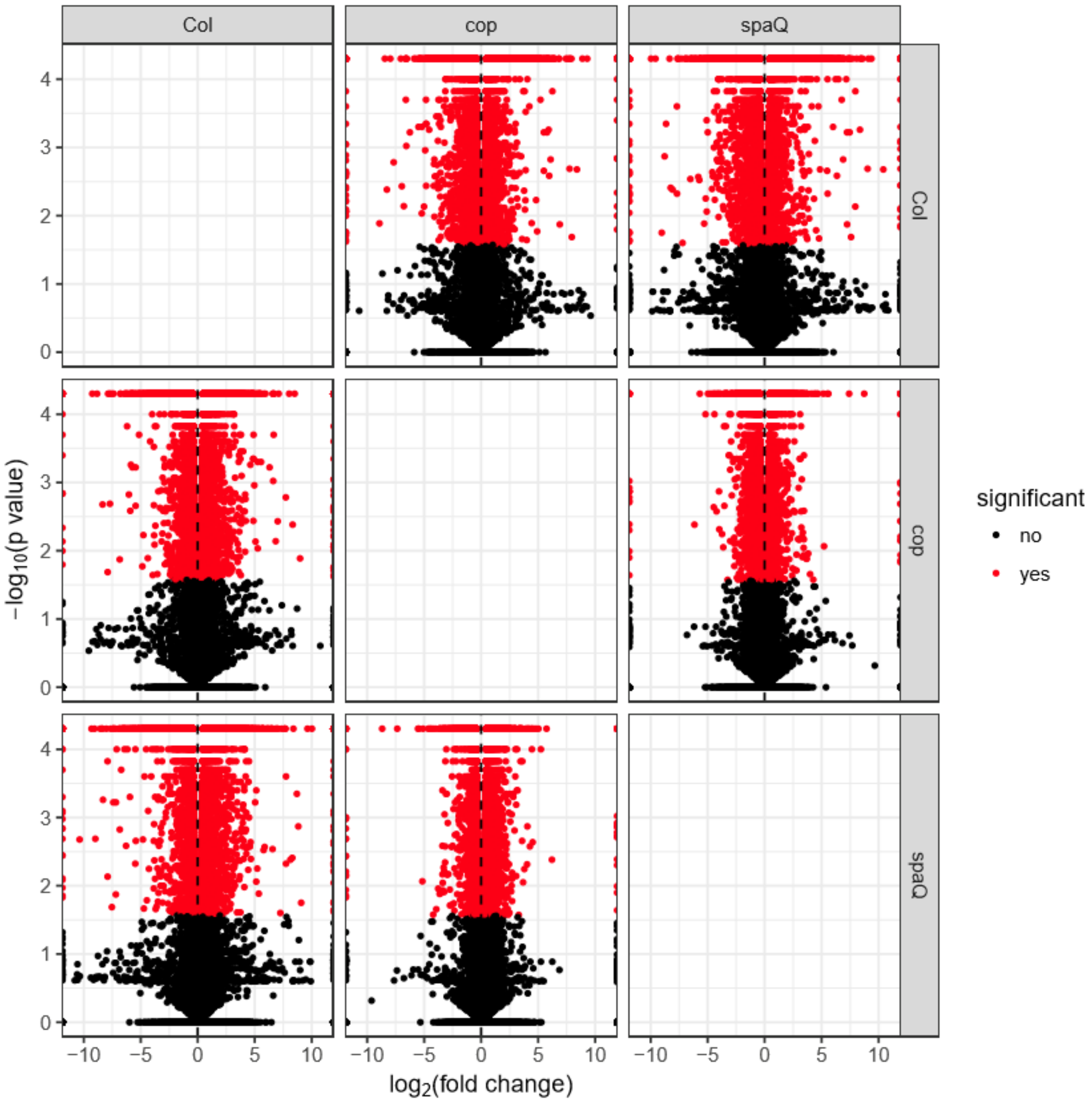


**Figure S4. Volcano plots for visualizing significantly differentially expressed genes in wild type (Col), *cop1-4* and *spaQ* mutants using Cufflinks and CummeRbund.**

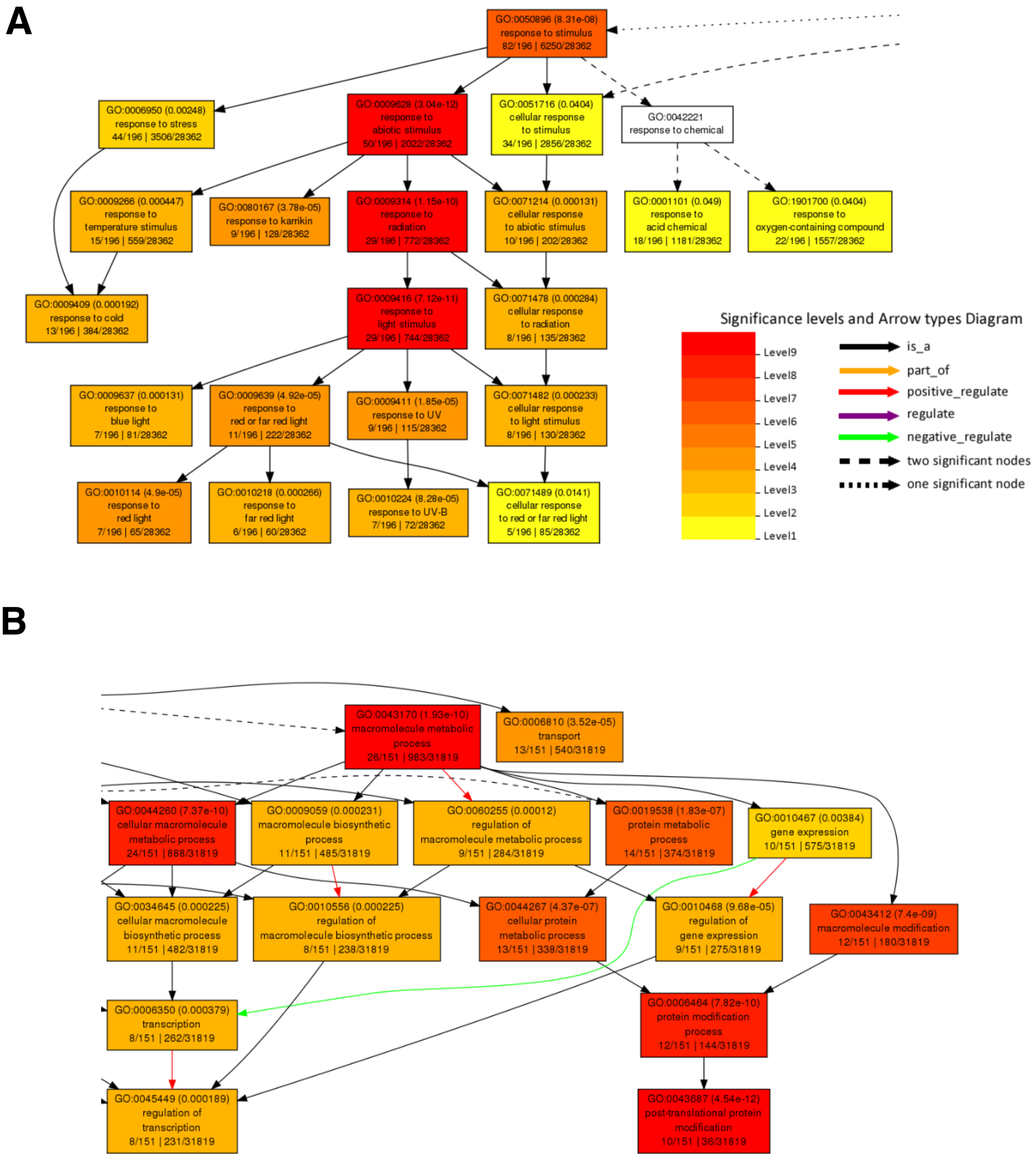


**Figure S5. Functional classification of SPA-regulated genes**

(A) Top 200 genes up-regulated in *spaQ* are involved in photosynthesis, response to light stimulus including far red, red and UV-B light. GO analysis were performed using agriGO -GO Analysis Toolkit and Database for Agricultural Community.

(B) Top 200 genes down-regulated in *spaQ* are related to response to auxin stimulus, post-translational protein modification, regulation of transcription. GO analysis were performed using agriGO -GO Analysis Toolkit and Database for Agricultural Community.

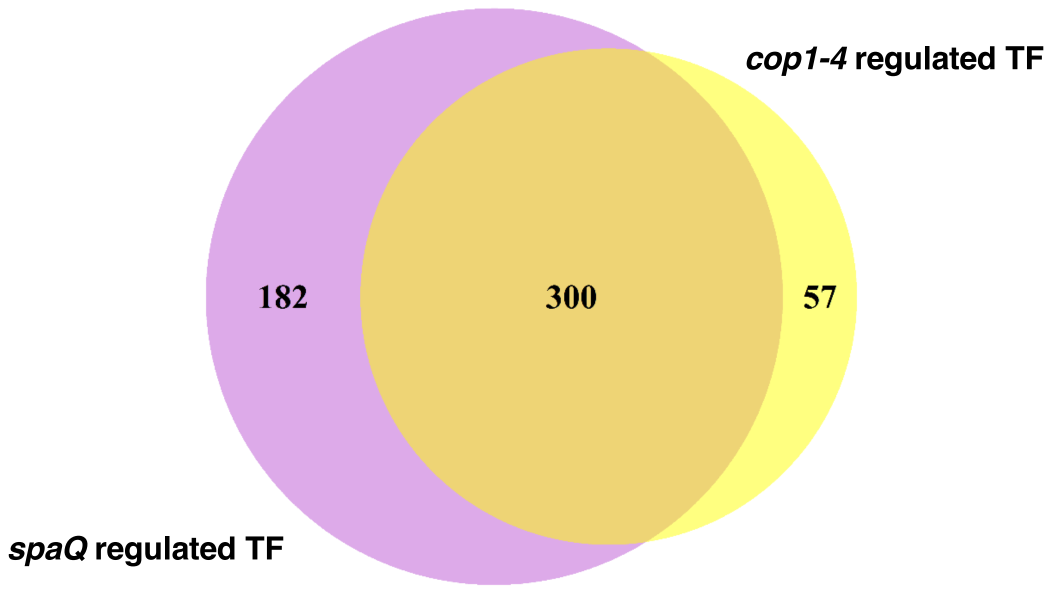


**Figure S6. Venn diagram shows the comparison between Transcription factor (TF) regulated in *cop1-4* and *spaQ* mutants.** Detail list of the TF direct target of SPAs independent of COP1 in Dataset S2.

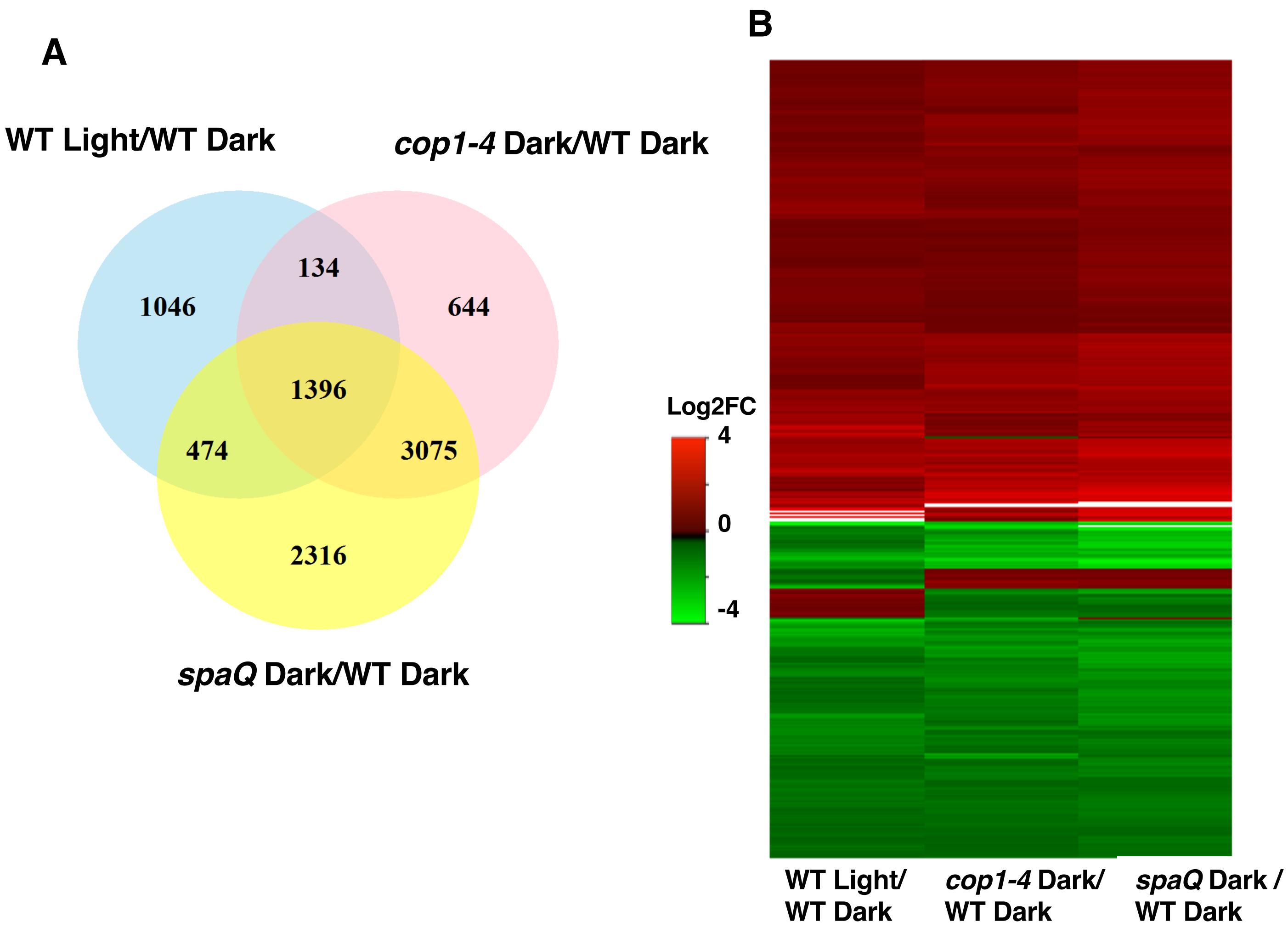


**Figure S7: Comparison of gene expression profiles between light-grown wild type seedlings, *cop1-4, and spaQ* in the dark.** Light regulated genes in wild type were obtained from Dong et al., 2014. wild type seeds were grown under 4 days in the dark and treated 6hrs of white light.

(A) Venn diagram shows the comparison of differential genes in WT light-grown seedlings, dark-grown seedling *cop1-4,* and *spaQ*. 1396 genes co-regulated by light-grown wild type and *cop1-4, spaQ* are presented.

(B) Heatmap shows the clustering expression patterns of 1396 genes co-regulated by light-grown WT, dark-grown *cop1-4* and dark-grown *spaQ.* WT Light/ WT Dark: expression ratios of white-light and dark-grown wild type seedlings, *cop1-4* Dark/ WT Dark: expression ratios of *cop1-4* dark grown and wild type dark-grown seedlings, *spaQ* Dark/ WT Dark: expression ratios of *spaQ* dark-grown and wild type dark-grown seedlings

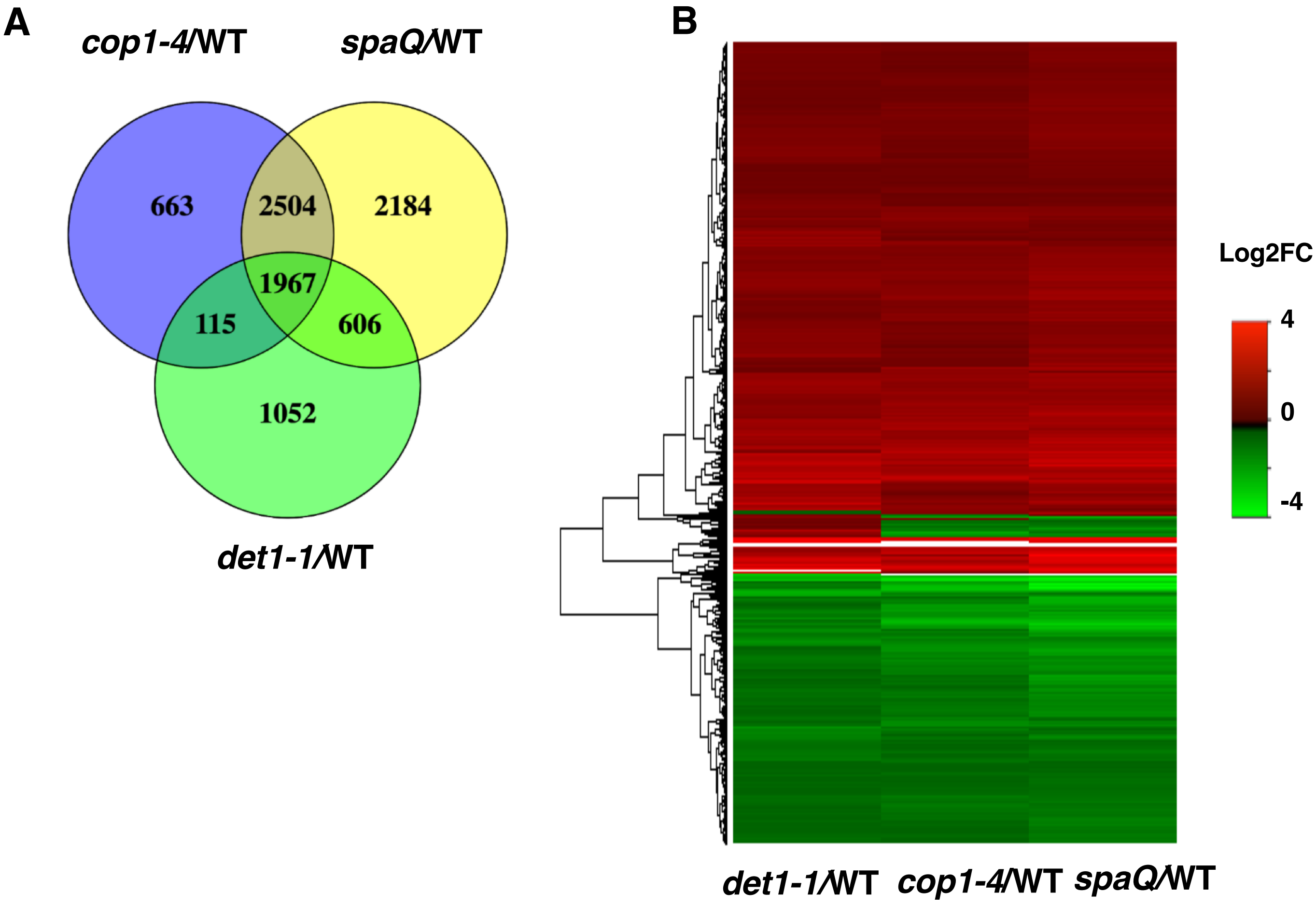


**Figure S8: Genomic expression profile comparison between dark-grown *cop1-4*, *spaQ,* and *det1-1* seedlings.** (A-B) Venn diagram (A) and Hierarchical clustering (B) of 1967 differentially expressed genes (DEG) in three different pairwise comparisons indicated (*cop1-4*/WT, *spaQ*/WT and *det1-1*/WT). Gene shown differential expression in three sample pairs are presented (fold change >2, FDR < 0.05). Color scale displays log2 fold-change value of significantly differential expression. Expression ratios were calculated between dark-grown mutant seedlings and wild-type dark-grown seedlings.

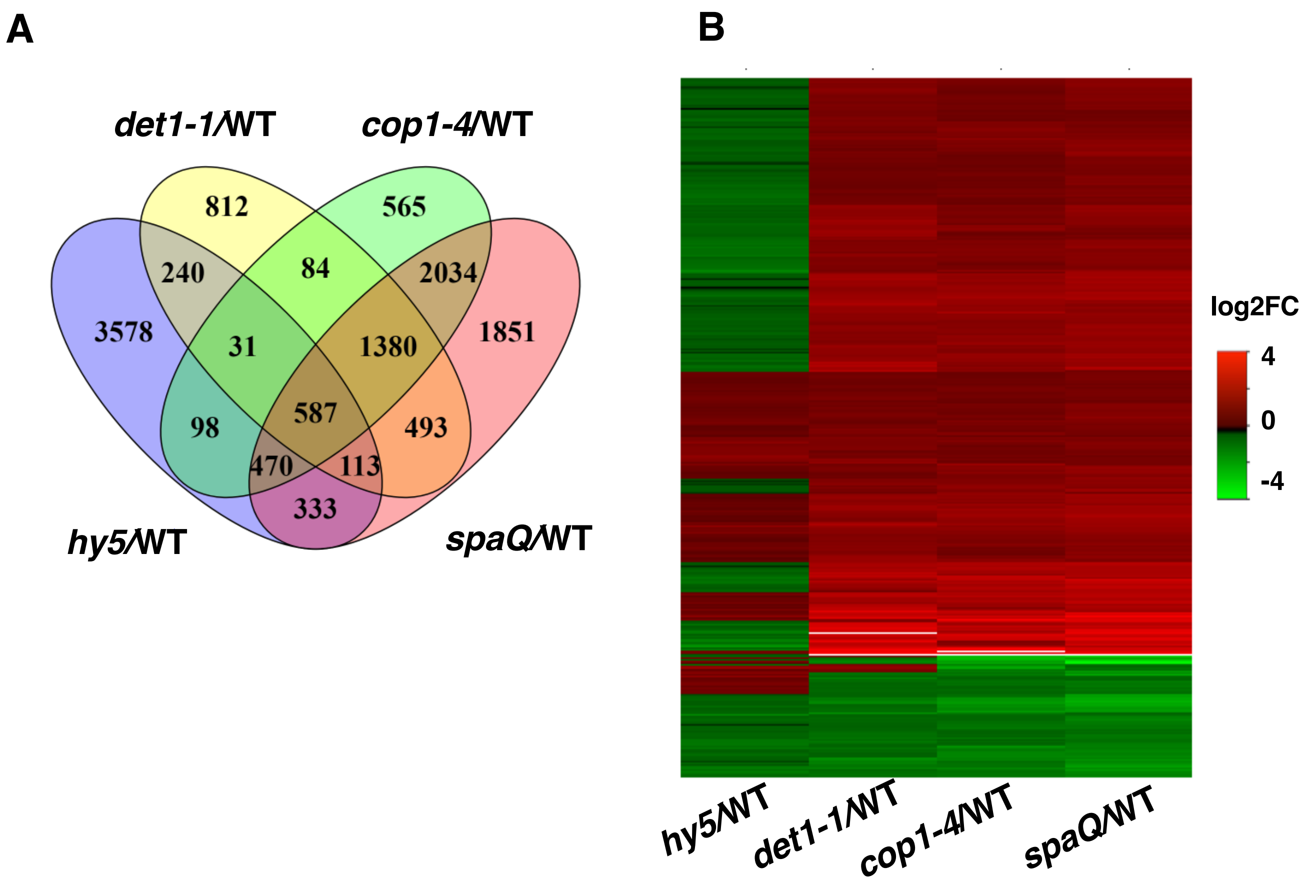


**Figure S9: Comparison of gene expression profiles between *cop1-4, spaQ, det1-1,* and *hy5* mutants.**

(A) Venn diagram shows the comparison of *hy5* differential genes with *cop1-4, spaQ* and *det1-1* regulated genes. 587 genes co-regulated by *cop1-4, spaQ, det1-1* and *hy5* mutants are shown.

(B) Heatmap shows the clustering expression patterns of 587 genes co-regulated by *cop1-4*, *spaQ, det1-1, and hy5 .* Expression ratios were calculated between mutant dark-grown seedlings and wild-type dark-grown seedlings.

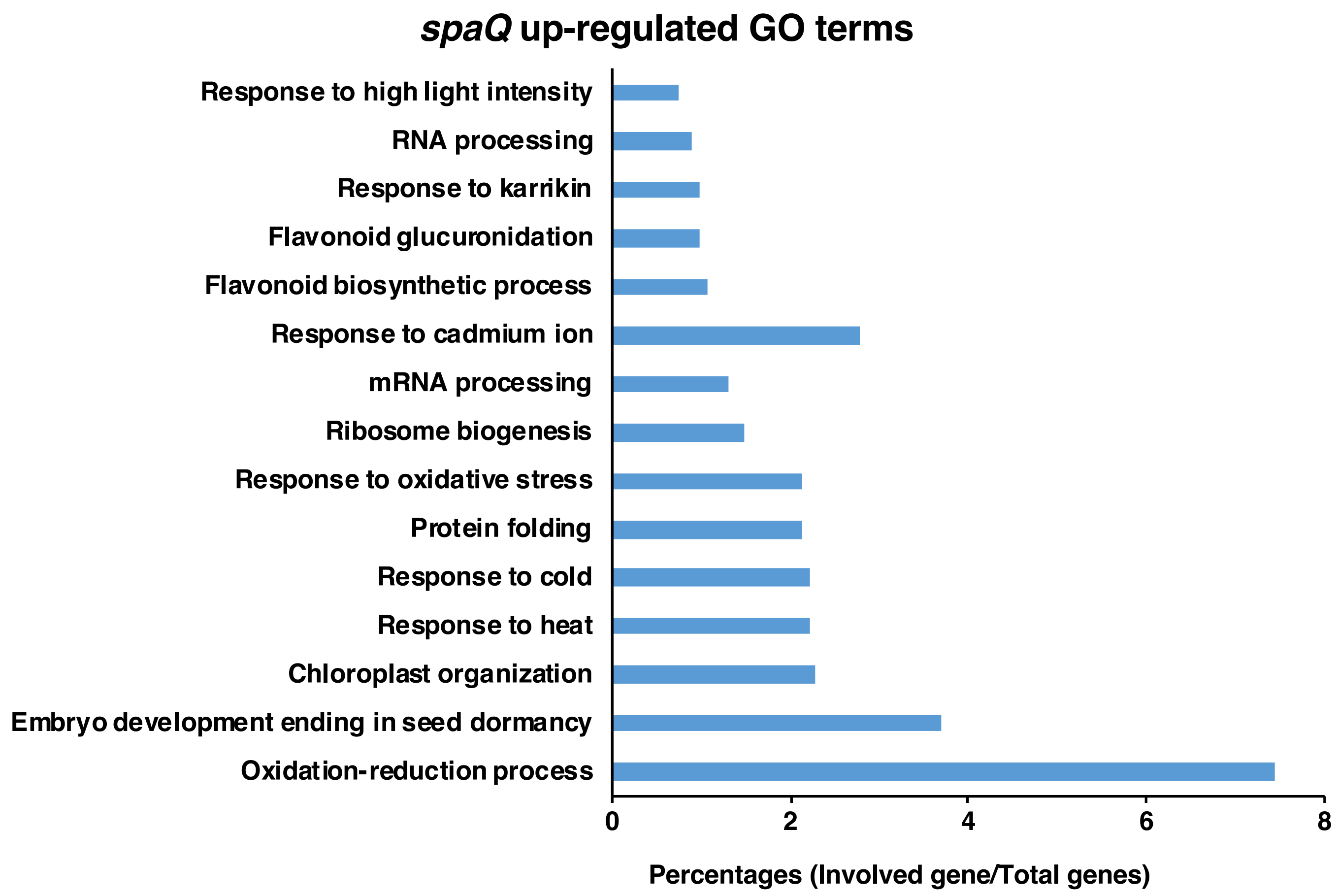


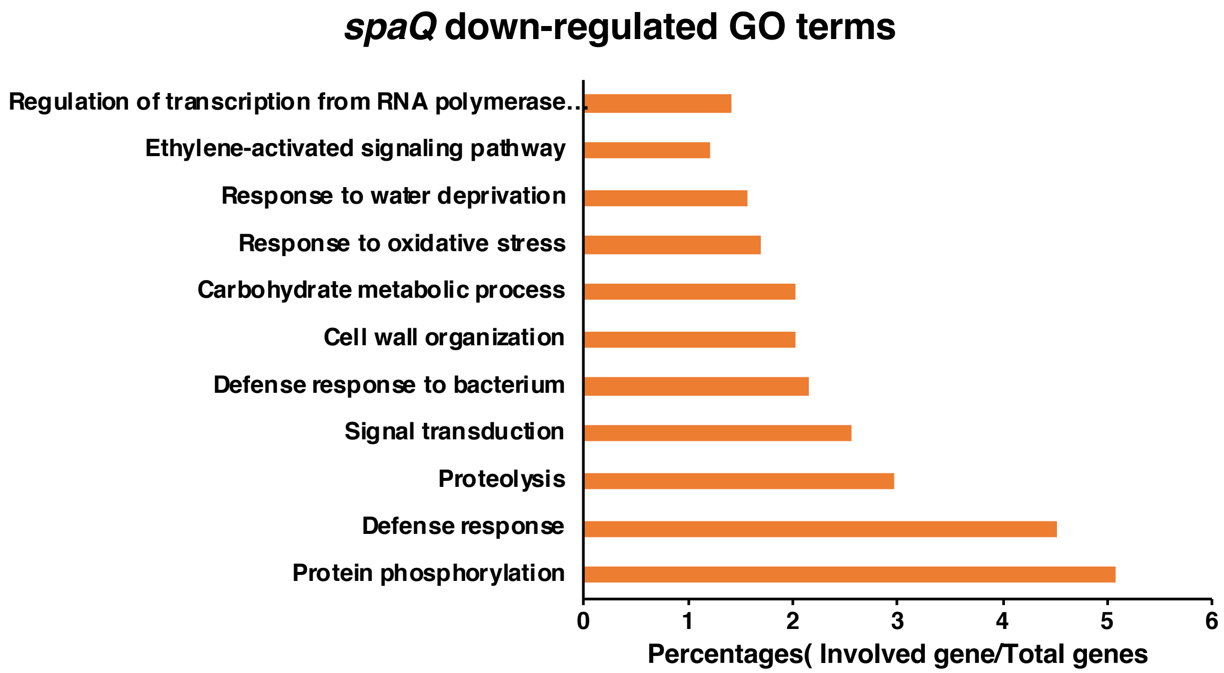


**Figure S10. GO analysis of 2790 genes which are differentially expressed in *spaQ* mutant but not in *cop1-4* mutant**. Only GO terms with p-values < 0.05 have been selected. GO analysis were performed using Database for Annotation, Visualization and Integrated Discovery (DAVID ).

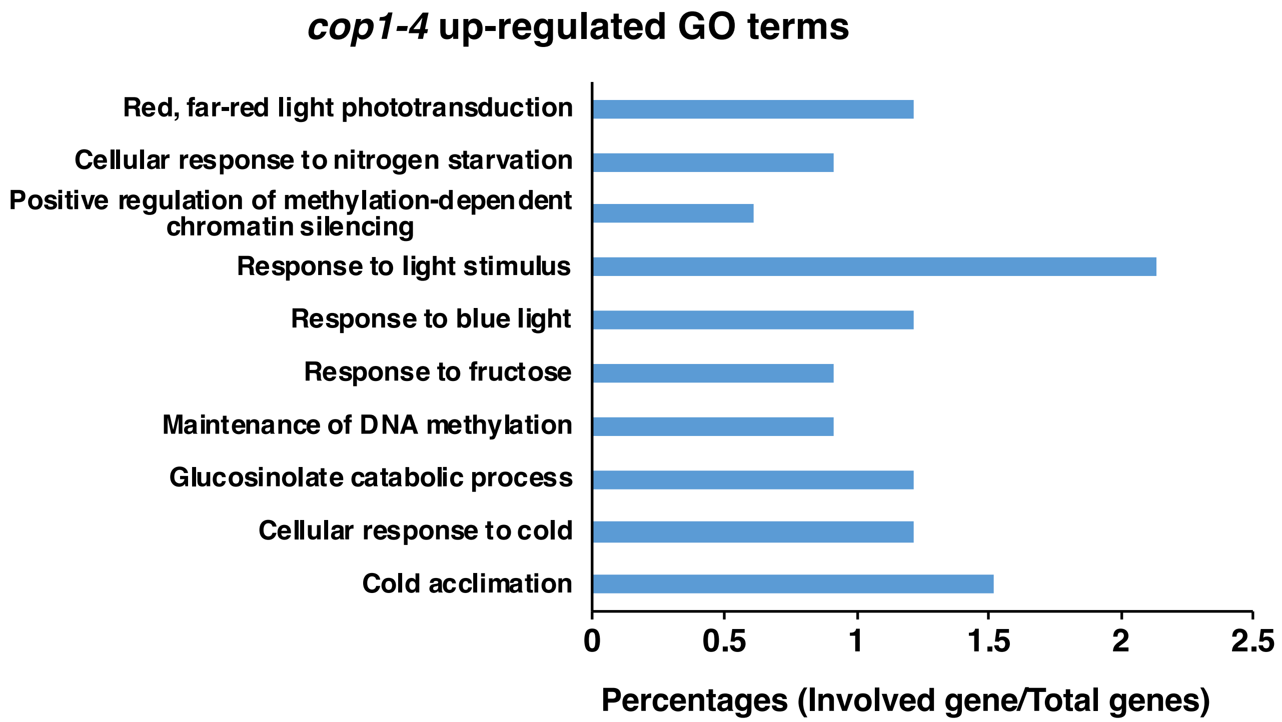


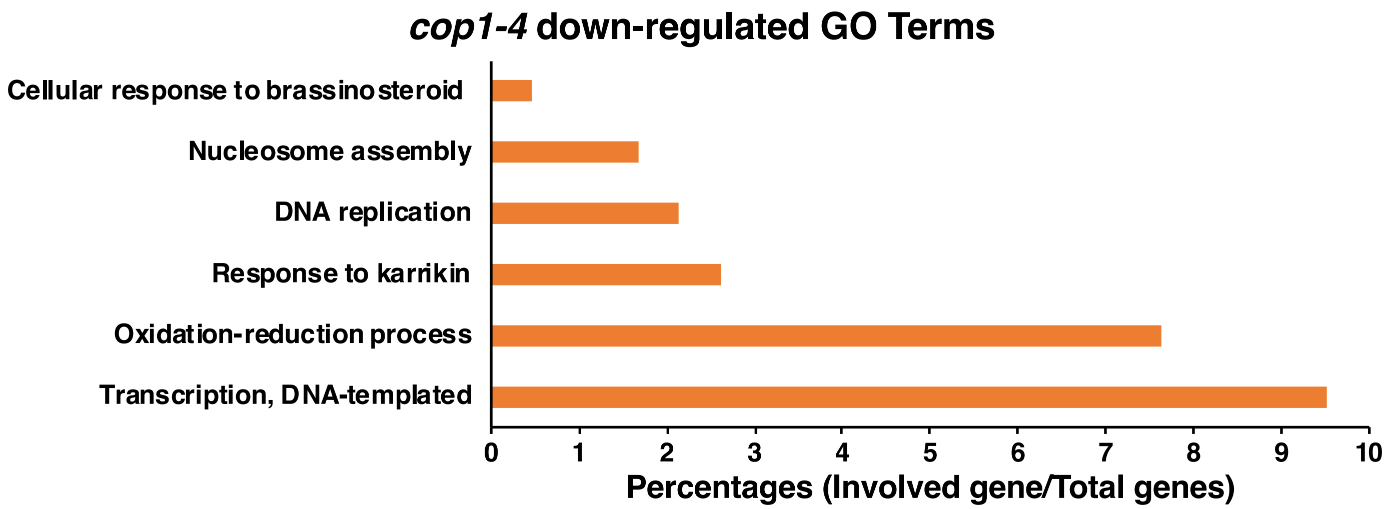


**Figure S11. GO analysis of 778 genes which are differentially expressed only in *cop1-4* mutant but not in *spaQ*.** Only GO terms with p-values < 0.05 have been selected. GO analysis were performed using Database for Annotation, Visualization and Integrated Discovery (DAVID ).

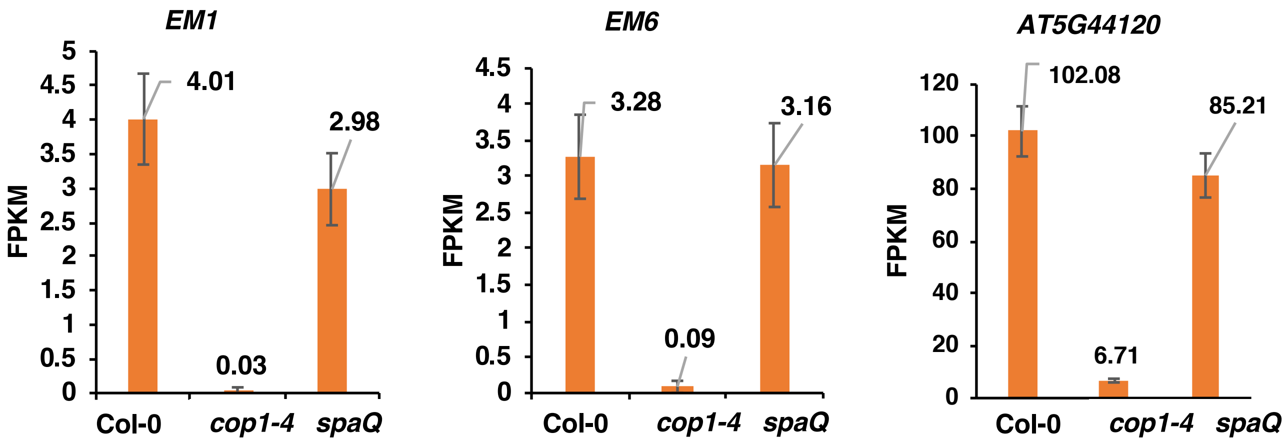


**Figure S12: The normalized expression level (FPKM; fragments per kilobase per million reads) of some differential expressed genes in *cop1-4* mutant, compared to wild-type (Col-0) and *spaQ* from RNA-seq data**

| **Table S1: RNA-seq reads and mapping statistics** | | | | | |
|---|---|---|---|---|---|
| **Sample** | **Total Reads** | **Read Mapped** | **Percent Mapped** | **Uniquely Mapped to a Single Gene** | **Percent Uniquely Mapped to a Single Gene** |
| Col-1 | 39116831 | 38377379 | 98.11 | 34739280 | 86.91 |
| Col-2 | 34130209 | 33475713 | 98.08 | 29859939 | 85.57 |
| Col-3 | 33606111 | 32942717 | 98.03 | 30330364 | 88.27 |
| *cop-1* | 42812777 | 41932304 | 97.94 | 36284077 | 82.69 |
| *cop-2* | 33652765 | 33101603 | 98.36 | 29147900 | 84.97 |
| *cop-3* | 27682983 | 27199292 | 98.25 | 24222174 | 85.75 |
| *spaQ-1* | 36771672 | 36232266 | 98.53 | 32156720 | 85.98 |
| *spaQ-2* | 45760739 | 45060737 | 98.47 | 41348272 | 88.82 |
| *spaQ-3* | 53400143 | 52799131 | 98.87 | 44487007 | 83.30 |
| Total | 346934230 | 341121142 | 98.29 | 297363657 | 85.81 |

| **Table S2. List of primers** | | | |
|---|---|---|---|
| **Gene** | **Forward Primer** | **Reverse Primer** | **Accession number** |
| CHS | AGCTGATGGACCTGCAGGCATCTTGGC | TGCATGTGACGTTTCCGAATTGTCGAC | AT5G13930 |
| CAB3 | GAGCTCAAGAACGGAAGATTGGC | CCGGGAACAAAGTTGGTTGC | AT1G29910 |
| ELIP2 | TCAACGGGAGACTAGCAATG | CCGTCAGAGATCTGAGCAAA | AT4G14690 |
| RBCS1A | ACCTTCTCCGCAACAAGTGG | GAAGCTTGGTGGCTTGTAGG | AT1G67090 |
| PER59 | TCTTCAGCAACCGTCTGTTC | TCCGTAGTGCTCCTGTCAAG | AT5G19890 |
| PRP2 | GTGTCCACCAAAGATTGCAC | ACTGGAGGCTTGTAGATGGG | AT2G21140 |
| HSP90.1 | CAC TAG GGA TGT GGA TGG GGA AC | CAC CTT CGT TTT CTT TCT TTG GTT C | AT5G52640 |
| HSP70 | ATCGACGCCAATGGTATCCT | AGAGCGTTCTTTGCATCCAC | AT3G12580 |
| GASA5 | AAATGGCGAATTGTATCAGAAG | TTAAGGACATTTTGGACGGC | AT3G02885 |
| KIN1 | ACCAACAAGAATGCCTTCCA | CCGCATCCGATACACTCTTT | AT5G15960 |
